# Hyperosmolarity-induced suppression of group B1 Raf-like protein kinases modulates drought-growth trade-off in *Arabidopsis*

**DOI:** 10.1101/2023.06.23.546348

**Authors:** Yoshiaki Kamiyama, Sotaro Katagiri, Kota Yamashita, Yangdan Li, Hinano Takase, Taishi Umezawa

## Abstract

When plants are exposed to drought stress, there is a trade-off between plant growth and stress responses. Here, we identified a signaling mechanism for the initial steps of the drought-growth trade-off. Phosphoproteomic profiling revealed that Raf13, a B1 subgroup Raf-like kinase, is dephosphorylated under drought conditions. Raf13 and the related B1-Raf Raf15 are required for growth rather than the acquisition of osmotolerance. We also found that Raf13 interacts with B55-family regulatory subunits of protein phosphatase 2A (PP2A), which mediates hyperosmolarity-induced dephosphorylation of Raf13. In addition, Raf13 interacts with an AGC kinase INCOMPLETE ROOT HAIR ELONGATION HOMOLOG 1 (IREH1), and Raf13 and IREH1 have similar functions in regulating cellular responses that promote plant growth. Overall, our results support a model in which Raf13-IREH1 activity promotes growth under nonstressed conditions, whereas PP2A activity suppresses Raf13-IREH1 during osmotic stress to modulate the physiological “trade-off” between plant growth and stress responses.

## Introduction

Plants often face a physiological “trade-off” between growth and stress responses, i.e. they have to make decisions whether to grow or to redirect energy to stress responses to survive under fluctuating environments ^1, 2^. Osmotic stress, including drought or high salinity, directly impacts plant growth and development, and a phytohormone abscisic acid (ABA) is involved in the process.

Recently, several reciprocal negative feedback loops between growth and ABA/osmotic stress signaling have been elucidated. TARGET OF RAPAMYCIN (TOR) kinase promotes growth signaling in the absence of stress, and TOR suppresses ABA signaling via phospho-regulation of ABA receptors ^3^. In the presence of stress, SNF1-related protein kinase 2 (SnRK2) is activated and inhibits TOR function ^3, 4^. Another system is group C Raf-like protein kinases (C-Raf), which promote growth by negatively regulating ABA signaling through phospho-dependent activation of PP2C thereby inhibiting SnRK2 activity. In the presence of stress, ABA-activated SnRK2s directly phosphorylate and inhibit C-Raf functions, thus allowing for heightened ABA signaling to occur ^5, 6^. In both cases, SnRK2 triggers the responses, suggesting that SnRK2 is a key factor involved in the trade-off between growth and stress responses.

There are 80 mitogen-activated protein kinase kinase kinases (MAPKKKs/MAP3Ks) in *Arabidopsis*, and 48 members are categorized as Raf-like families and can further be grouped into 11 subgroups; B1 to B4 and C1 to C7 ^7^. Recent studies revealed the direct regulation of SnRK2s by group B Raf-like protein kinases. In *Physcomitrium patens*, a B3-Raf, *ABA AND ABIOTIC STRESS-RESPONSIVE RAF-LIKE KINASE* (*ARK*), is required for the SnRK2 activation, gene expression, and acquisition of osmotolerance when exposed to osmotic stress ^8^. *Arabidopsis* B2/B3-Raf and B4-Raf also regulate subclass III SnRK2s and subclass I SnRK2s, respectively, in response to hyperosmolarity ^9–13^. However, despite the growing evidence of group B Rafs in SnRK2 regulation under osmotic stress conditions, the physiological roles or the signaling factors associated with B1-Rafs are still unknown.

Here, we found that a B1-Raf Raf13 is dephosphorylated during osmotic stress responses to lead to its inactivation and destabilization. B1-Raf consists of Raf13, Raf14 and Raf15 in Arabidopsis, and Raf13 and Raf15 function redundantly in promoting plant growth under optimal growth conditions. Furthermore, we demonstrated that Raf13/Raf15 function in concert by interacting with a protein kinase IREH1 and protein phosphatase PP2A holoenzyme(s) containing a B55-family regulatory subunit. These results reveal that the B1-Raf/IREH1/PP2A kinase-phosphatase complex provides an SnRK2-independent mechanism of modulating the “trade-off” between plant growth and stress responses.

## Results

### Raf13 is dephosphorylated in response to osmotic stress

Protein phosphorylation/dephosphorylation has significant roles in drought stress signaling in plants^14–16^. We performed a label-free quantitative phosphoproteomic analysis of Arabidopsis wild-type seedlings under drought treatment for 0, 3, 5, and 9 days. The result showed that several phosphopeptides from B3-Raf (Raf4/AtARK1), B4-Rafs (Raf18 and Raf20), SnRK2s (SRK2B, SRK2D, SRK2E and SRK2I), or SnRK2 substrates (VARICOSE, AREB/ABFs, SLAC1, AKS1 and FREE1), were upregulated, consistent with previous reports ^10, 13, 17–22^ (Fig. 1a,b and Supplementary Table 1). We focused on a phosphopeptide from the B1-Raf Raf13. The peptide is phosphorylated on two serine residues (Ser-310 and Ser-313, KLEGYPNA*p*SGS*p*SLR), and was significantly downregulated at 5 and 9 days (Fig. 1b and Supplementary Table 1). This suggested that Raf13 could be dephosphorylated in response to drought stress. This result was further confirmed by immunoblotting of protein extracts from Arabidopsis transgenic plants expressing *Raf13-CFP-HA* or *AtARK1-HA*. Raf13-CFP-HA showed electrophoretic mobility shifts with lower molecular mass after dehydration treatment for 30 min, whereas AtARK1-HA protein shifted to higher molecular mass (Fig. 1c). Such mobility shifts were not observed during ABA treatment (Extended Data Fig. 1), suggesting that Raf13 is specifically dephosphorylated in response to osmotic stress. An additional LC-MS/MS analysis from wild-type seedlings treated with or without dehydration stress detected six peptides containing nine phosphosites (Ser-10, Ser-38, Ser-310, Ser-313, Ser-335, Ser-390, Ser-465, Ser-468, and Ser-469) in Raf13 (Extended Data Fig. 2), and four phosphopeptides containing six phosphosites (Ser-38, Ser-310, Ser-313, Ser-335, Ser-468, and Ser-469) showed a significant decrease after dehydration for 30 min (Fig. 1d,e and Supplementary Table 2). These results indicated that Raf13 is multiply dephosphorylated *in planta* during osmotic stress responses.

**Fig. 1.**
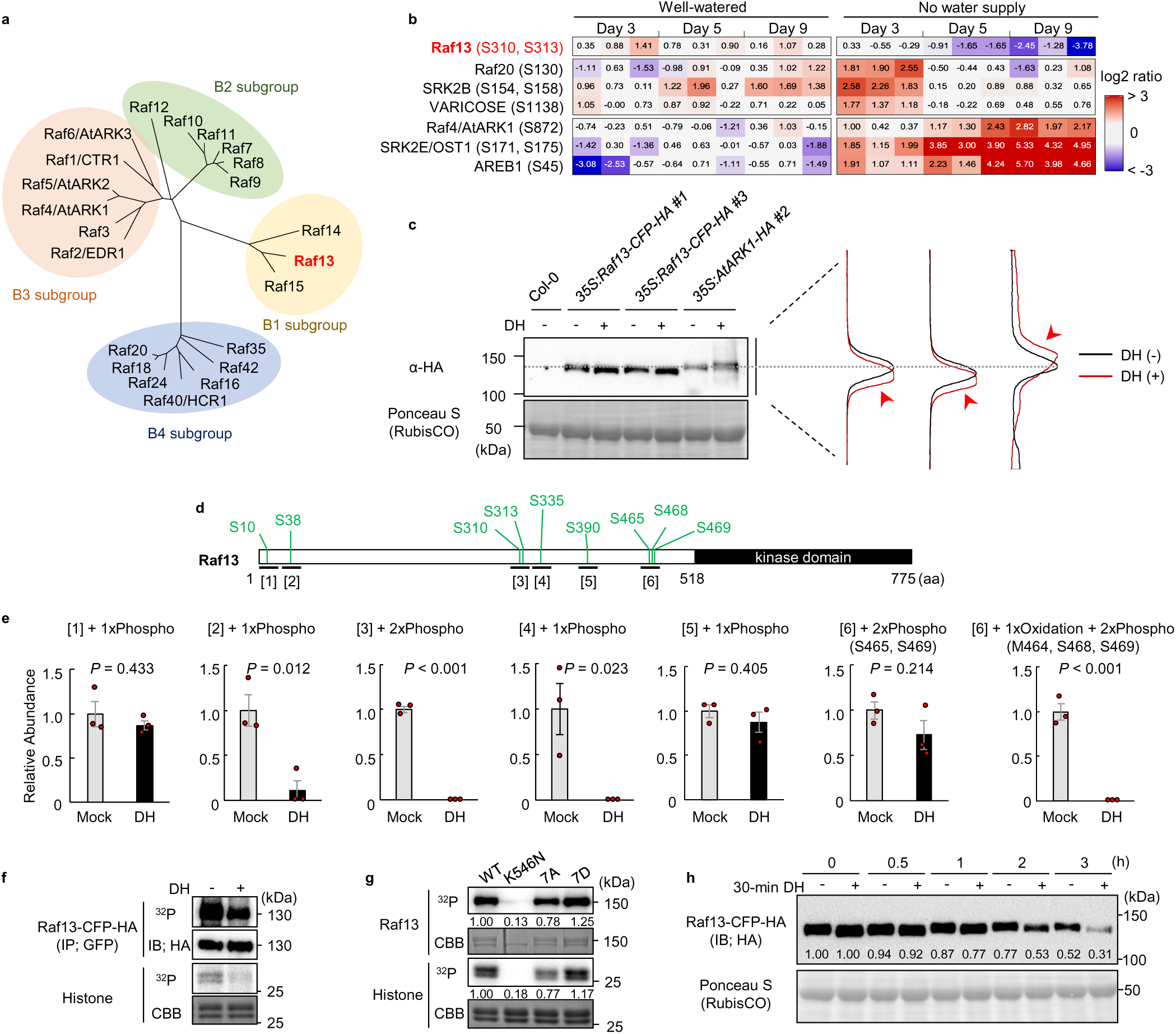
Raf13 is dephosphorylated and suppressed in response to hyperosmolarity. **a**, Phylogenetic tree of *Arabidopsis* group B Raf-like protein kinases. The tree was generated based on the putative kinase catalytic domains. **b,** Heat map showing the relative phosphopeptide abundances of group B-Rafs, SnRK2s, or SnRK2 substrates in the aerial parts of soil-grown *Arabidopsis* wild-type (Col-0) seedlings with/without water supply for the indicated time periods. **c,** Western blotting showing the band shifts of Raf13-CFP-HA or AtARK1-HA with/without dehydration (DH) for 30 min. Intensity profiles of each Raf13-CFP-HA or AtARK1-HA were measured and shown in the panel by normalizing the peak value to 1 (left: *35S:Raf13-CFP-HA #1*, middle: *35S:Raf13-CFP-HA #*3, right: *35S:AtARK1-HA #2*). Red arrowheads indicate electrophoretic mobility shifts. **d,e,** LC-MS/MS analysis identified phosphopeptides ([1]-[6]) derived from Raf13 protein (**d**) and their relative abundance (**e**) in *Arabidopsis* wild-type seedlings with/without DH treatment for 30 min. Data are means ± SE (n =3), and significance was determined using two-tailed Student’s *t-*test. **f,** IP-kinase assay showing Raf13 protein kinase activity from *Arabidopsis 35S:Raf13-CFP-HA* seedlings with/without DH treatment for 30 min. **g,** *In vitro* phosphorylation assay showing protein kinase activities of MBP-tagged Raf13 variants purified from *E. coli*. Kinase-dead Raf13 (K546N) was used as a negative control. Values are means of signal intensities from auto-and histone-phosphorylation measured and normalized to that of WT (n=3). **h,** *In vitro* cell-free degradation assay showing the protein stability of phosphorylated- or dephosphorylated-Raf13 protein. Crude extracts from *35S:Raf13-CFP-HA* seedlings treated with/without DH for 30 min were incubated at 21 °C for indicated periods, and values are means of Raf13-CFP-HA protein levels measured and normalized to that at 0 h (n=3).

**Fig. 2.**
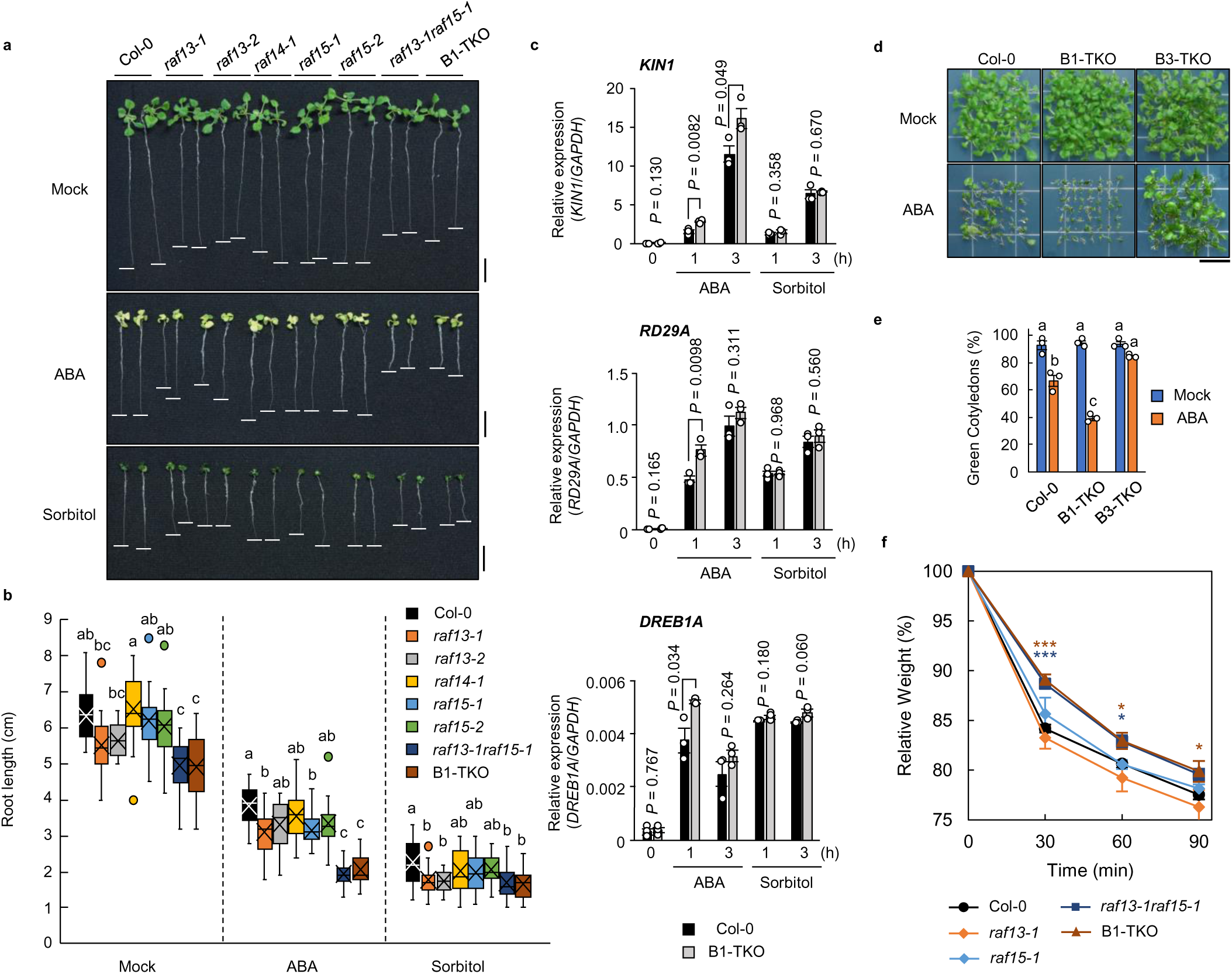
B1-Raf mutants show reduced growth and hypersensitivity to ABA. **a,b**, Primary root length of 11-day-old Arabidopsis wild-type (Col-0) and B1-Raf mutants under 10 μM ABA or 400 mM sorbitol. Scale bars, 1 cm. Data are presented as boxplot; box limits represent the first and the third quartiles with the medians and the means marked as horizontal lines and cross marks, respectively. The maximum length of the whiskers is 1.5 times the interquartile range (IQR) from the first and the third quartiles, and points outside this range are shown as outliers. Different letters indicate significant differences (Tukey’s test, *P* < 0.01, n = 24). **c,** Relative expression levels of ABA- or osmotic stress-responsive genes in 2-week-old Arabidopsis wild-type and B1-TKO seedlings treated with 50 μM ABA or 400 mM sorbitol for the indicated time periods. *GAPDH* was used as an internal control. Bars indicate means ± SE (n = 3), and significance was determined using two-tailed Student’s *t-*test. **d,** Cotyledon greening of wild-type, B1-TKO and B3-TKO with/without 0.5 μM ABA for 9 days. **e,** Seedlings showing green cotyledons at 5 days in **d** were counted, and data are presented as means ± SE (n = 3). Different letters indicate significant differences (Tukey’s test, *P* < 0.05). Each replicate contains 36 seeds. Scale bar, 1 cm. **f,** Water loss from detached leaves of wild-type and B1-Raf mutants. Data are means ± SD (n = 3), and asterisks indicate significant differences by Dunnett’s test (\**P* < 0.05, \*\*\**P* < 0.001). Each replicate consists of five individual leaves.

We next used immunoprecipitation-kinase (IP-kinase) assays to investigate possible functions of Raf13 dephosphorylation. As shown in Fig. 1f, autophosphorylation and histone transphosphorylation activities of Raf13-CFP-HA were reduced when Raf13-CFP-HA was immunoprecipitated from proteins extracted from seedlings treated with dehydration stress for 30 min. Consistent with these results, the kinase activities of recombinant maltose-binding protein (MBP)-tagged Raf13 (MBP-Raf13) were enhanced or reduced by Asp or Ala substitutions, respectively, at seven Ser residues that correspond to detected phosphorylation sites, including Ser-38, Ser-310, Ser-313, Ser-335, Ser-465, Ser-468, and Ser-469 (Fig. 1g). To assess if phosphorylation status impacts Raf13 stability, we conducted an *in vitro* cell-free degradation assay. Crude extracts were prepared from *35S:Raf13-CFP-HA* seedlings treated with or without 30-min-dehydration stress, and the extracts were incubated at 21°C for indicated periods. As shown in Fig. 1h, the protein abundances of both phosphorylated- and dephosphorylated-Raf13 were decreased in a time-dependent manner. However, a much faster decrease in abundance was observed for the dephosphorylated form. Together, these data suggested that hyperosmolarity-induced dephosphorylation suppresses Raf13 kinase catalytic activity and protein stability.

### B1-Rafs are required for plant growth rather than the acquisition of osmotolerance

To gain insight into the physiological roles of three B1-Rafs, we next obtained mutants with individual transfer DNA (T-DNA) insertions in *Raf13*, *Raf14* and *Raf15* (Extended Data Fig. 3a), and generated a double mutant (*raf13-1raf15-1*) and a triple mutant (*raf13-1raf14-1raf15-1*; B1-TKO). These mutants showed reduced growth relative to wild-type under optimal growth conditions. In this regard, significantly shorter primary roots were observed in *raf13-1raf15-1* and B1-TKO compared with wild-type under control conditions. Primary roots were slightly shortened in *raf13-1* and *raf13-2*, but not in *raf14-1*, *raf15-1* or *raf15-2* (Fig. 2a,b). In the presence of ABA, primary root elongation was inhibited by 39.8% in the wild-type and by approximately 60% in the double/triple mutants. On the other hand, 400 mM sorbitol inhibited root elongation similarly in wild-type and mutants (Fig. 2a,b). Some ABA- and stress-responsive genes, *KIN1*, *RD29A* and *DREB1A*, were hyper-induced in B1-TKO seedlings in response to 50 µM ABA treatment, but the expression level was similar between wild-type and B1-TKO in response to 400 mM sorbitol (Fig. 2c).

**Fig. 3.**
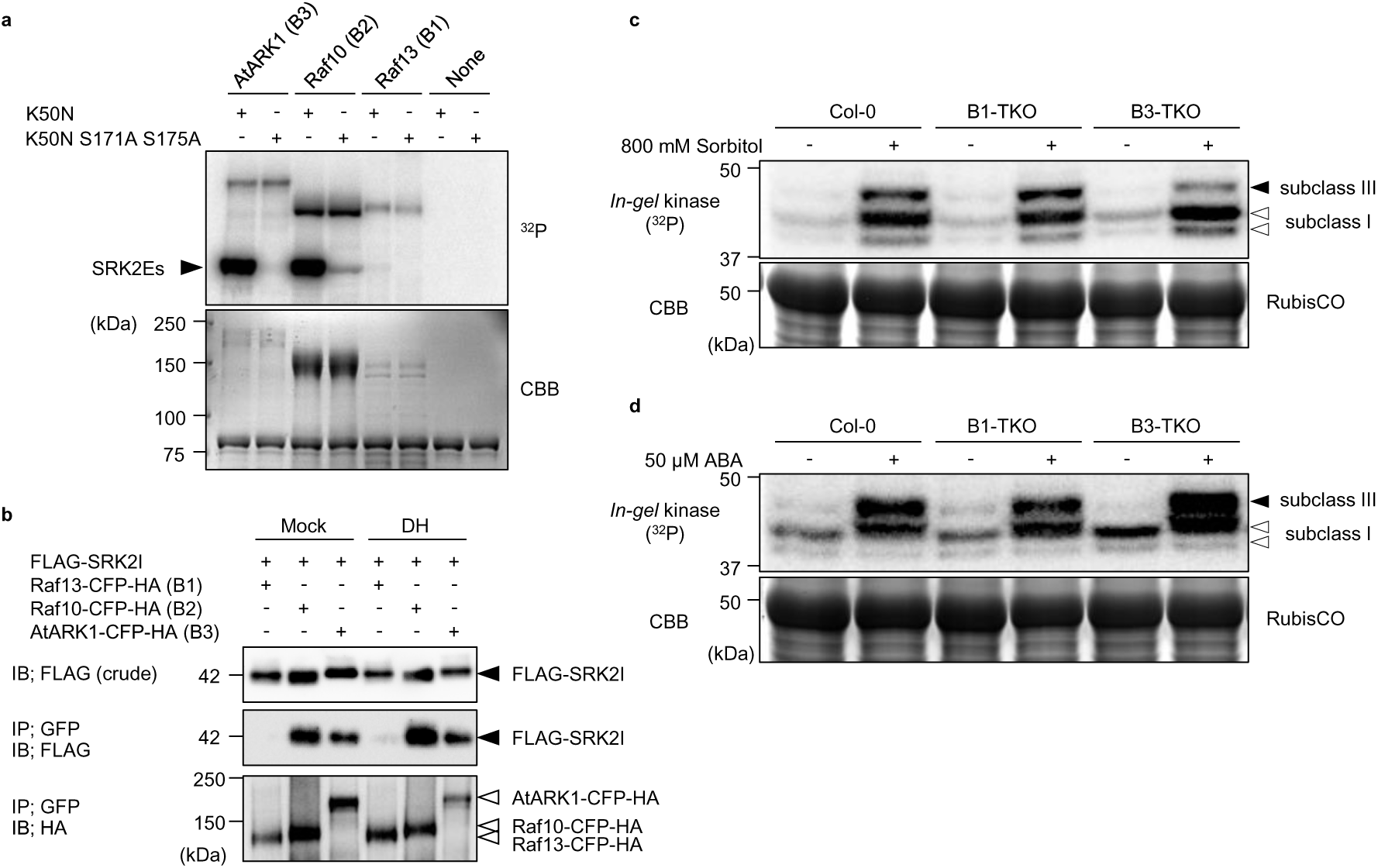
B1-Rafs function independently of SnRK2. **a**, *In vitro* phosphorylation assay of kinase-dead SRK2E (K50N) and SRK2E (K50N S171A S175A) proteins by Raf13. All recombinant proteins were purified as MBP fusions. **b,** Co-IP assay of Raf13 and SRK2I from *N. benthamiana* leaves treated with/without dehydration (DH) for 1 h. IP and IB indicate immunoprecipitation and immunoblotting, respectively. AtARK1/Raf4 (B3 subgroup) and Raf10 (B2 subgroup) in **a,b** were used as positive controls. **c,d,** *In-gel* kinase assays of proteins extracted from 2-week-old seedlings of wild-type (Col-0), B1-TKO and B3-TKO plants treated with 800 mM sorbitol (**c**) or 50 µM ABA (**d**) for 30 min using histone IIIS as substrates. Black arrows and open arrows indicate the position of subclass III and subclass I SnRK2s, respectively. Autoradiography (^32^P) and Coomassie brilliant blue (CBB) staining in **a,c,d** show phosphorylation and loading, respectively.

B1-TKO showed a significantly lower cotyledon greening rate relative to wild-type plants in the presence of ABA, but wild-type and B1-TKO were similar in the absence of ABA (Fig. 2d,e and Extended Data Fig. 3b,c). Conversely, *35S:Raf13-CFP-HA* plants displayed an ABA-insensitive phenotype (Extended Data Fig. 3d,e). Such a phenotype in B1-TKO is opposite to that of B3-Raf triple knockout mutant (raf4raf5raf6; B3-TKO) (Fig. 2d,e). No significant change in seed germination rates was observed in the mutants (Extended Data Fig. 3f). We further measured water loss from detached leaves, and found that the lack of both *Raf13* and *Raf15* genes resulted in reduced water loss (Fig. 2f), which is probably attributed to reduced stomatal aperture even before ABA treatment (Extended Data Fig. 3g). Notably, the β-glucuronidase (GUS) activity was observed in young leaves, guard cells, and roots in *Arabidopsis Raf13pro:GUS* plants (Extended Data Fig. 3h). Collectively, these results indicated that B1-Rafs are required for some of the aspects of plant growth and appear to negative regulate responses to exogenous ABA.

### B1-Rafs function independently of SnRK2 regulation

Phenotyping tests suggested that B1-Rafs might have different functions from B2-,B3-, and B4-Rafs, some of which are upstream activators of SnRK2s ^9–13^. To check the biochemical properties of Raf13, we prepared the recombinant MBP-tagged Raf13 with autophosphorylation and histone transphosphorylation activities. A K546N substitution in Raf13 abolished all kinase activity, indicating that Raf13 is a canonical protein kinase (Extended Data Fig. 4a). Raf4/AtARK1 (B3) and Raf10 (B2) strongly phosphorylated SRK2E (K50N) in the kinase activation loop (Fig. 3a). However, Raf13 did not phosphorylate SRK2E (Fig. 3a) and SRK2G (Extended Data Fig. 4b).

**Fig. 4.**
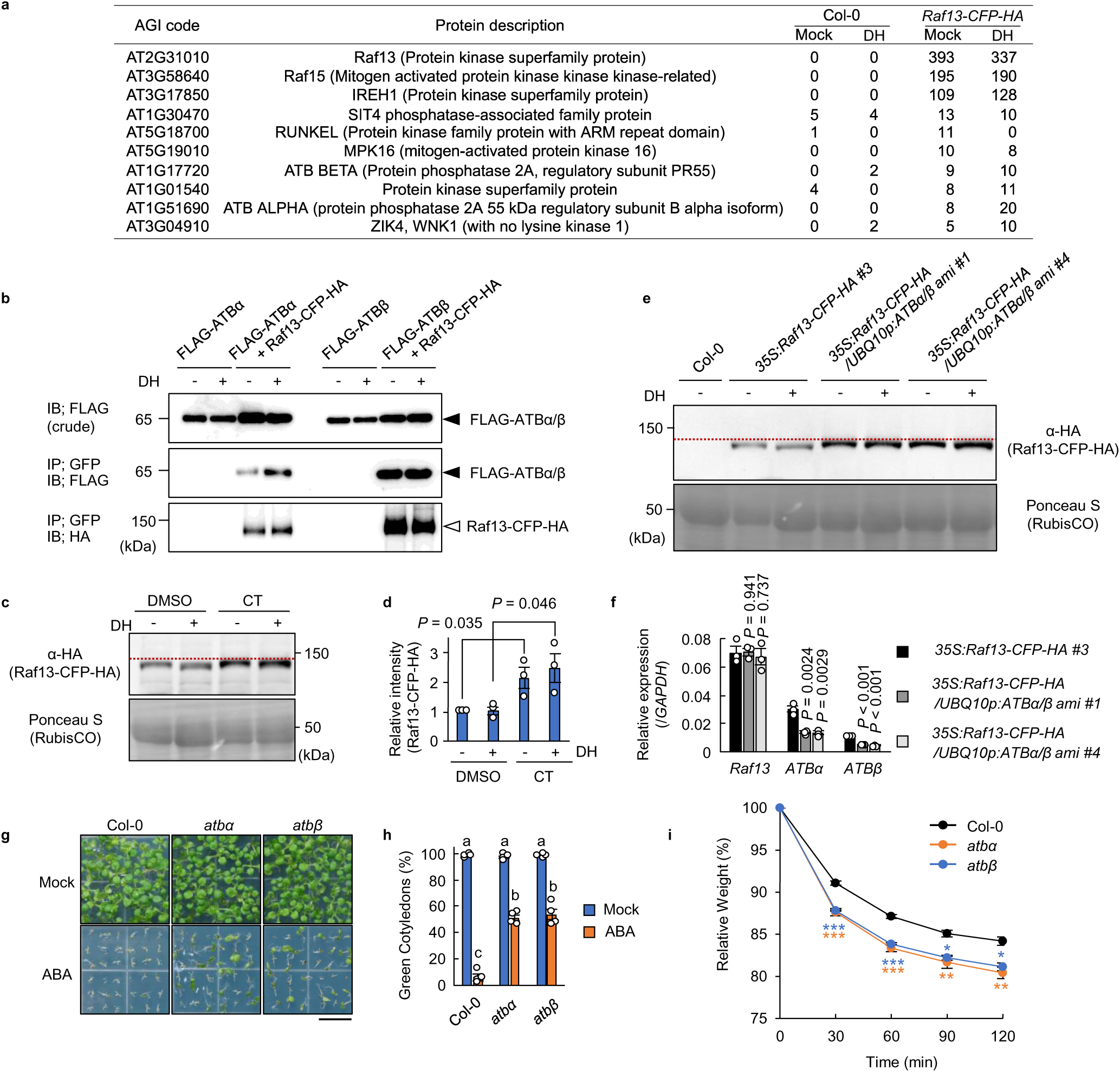
Osmotic stress-dependent Raf13 dephosphorylation is mediated by type 2A protein phosphatases. **a**, List of the protein kinases/phosphatases suggested to potentially interact with Raf13-CFP-HA protein from 2-week-old Arabidopsis seedlings treated with/without dehydration (DH) for 30 min. The sum of the peptide spectrum match (PSM) scores from the two independent biological replicates were shown. The cut-off was made with the PSM scores greater than or equal to 10 in *35S:Raf13-CFP-HA* in either Mock or DH conditions and less than 6 in wild-type at both Mock and DH conditions. **b,** Co-IP assays of Raf13 and ATBα/β from *N. benthamiana* leaves treated with/without DH for 1 h. **c,d,** Western blotting of Raf13-CFP-HA under cantharidin (CT) treatment (**c**) and relative amounts of Raf13-CFP-HA protein measured using ImageJ (**d**). Arabidopsis *35S:Raf13-CFP-HA* plants were incubated for 12 h under 50 μM CT or dimethyl sulfoxide (DMSO) control, and then the seedlings were treated with/without dehydration (DH) for 30 min. The red dotted line in **c** aligns the slower migrating forms in CT-treated plants. Data are means ± SE (n = 3), and significance was determined using two-tailed Student’s *t*-test (**d**). **e,f,** Western blotting of Raf13-CFP-HA in *ATBα/β*-knockdown plants treated with/without DH for 30 min (**e**), and quantification of *Raf13*, *ATBα* or *ATBβ* transcripts normalized to *GAPDH* by qRT-PCR (**f**). The red dotted line in **e** aligns the slower migrating forms in the *ATBα/β*-knockdown plants. Bars indicate means ± SE (n = 3), and significance was determined using two-tailed Student’s *t-*test (**f**). **g,h,** Cotyledon greening rates of wild-type, *atbα* or *atbβ* plants with/without 1 μM ABA for 8 days (**g**) and data are presented as means ± SE (n = 4) (**h**). Different letters in **h** indicate significant differences (Tukey’s test, *P* < 0.01). Each replicate contains 54 seeds. Scale bar, 1 cm. **i,** Water loss from detached leaves of wild-type, *atbα* and *atbβ* plants. Data are means ± SE (n = 3), and asterisks indicate significant differences by Dunnett’s test (\**P* < 0.05, \*\**P* < 0.01, \*\*\**P* < 0.001). Each replicate consists of five individual leaves.

Co-immunoprecipitation (Co-IP) assays showed no or negligible interaction between Raf13 and SRK2I, but both Raf4/AtARK1 and Raf10 interacted with SRK2I (Fig. 3b). Furthermore, *in-gel* kinase assays demonstrated that 800 mM sorbitol- or 50 µM ABA-induced SnRK2 activation was not affected in B1-TKO or *35S:Raf13-CFP-HA* plants, but 800 mM sorbitol-induced SnRK2 activation was reduced in B3-TKO as previously reported ^9^ (Fig. 3c,d and Extended Data Fig. 4c,d). Collectively, these results indicate that B1-Rafs function independently of SnRK2s.

### PP2A holoenzymes mediate Raf13 dephosphorylation in response to hyperosmolarity

To identify signaling factors associated with Raf13, we performed immunoprecipitation followed by LC-MS/MS analyses (IP-MS). Raf13-CFP-HA protein was immunoprecipitated from *Arabidopsis 35S:Raf13-CFP-HA* plants with or without dehydration treatment for 30 min. As a result, a total of 37 proteins were identified as Raf13-CFP-HA-interacting candidates under the control and/or dehydration conditions (Fig. 4a and Supplementary Table 3). In the dataset, we found two subunits of type 2A protein phosphatase (PP2A), ATBα and ATBβ, also known as B55α or B55β, respectively. To further confirm the physical interactions between Raf13 and ATBα/β, we performed Co-IP assays in *N. benthamiana* leaves, and found that FLAG-ATBα and -ATBβ were co-immunoprecipitated with Raf13-CFP-HA under both control and dehydration conditions (Fig. 4b). Raf13 and ATBα were localized in the cytosol, and ATBβ was localized in both cytosol and nuclei (Extended Data Fig. 5a), suggesting possible colocalization in the cytosol.

**Fig. 5.**
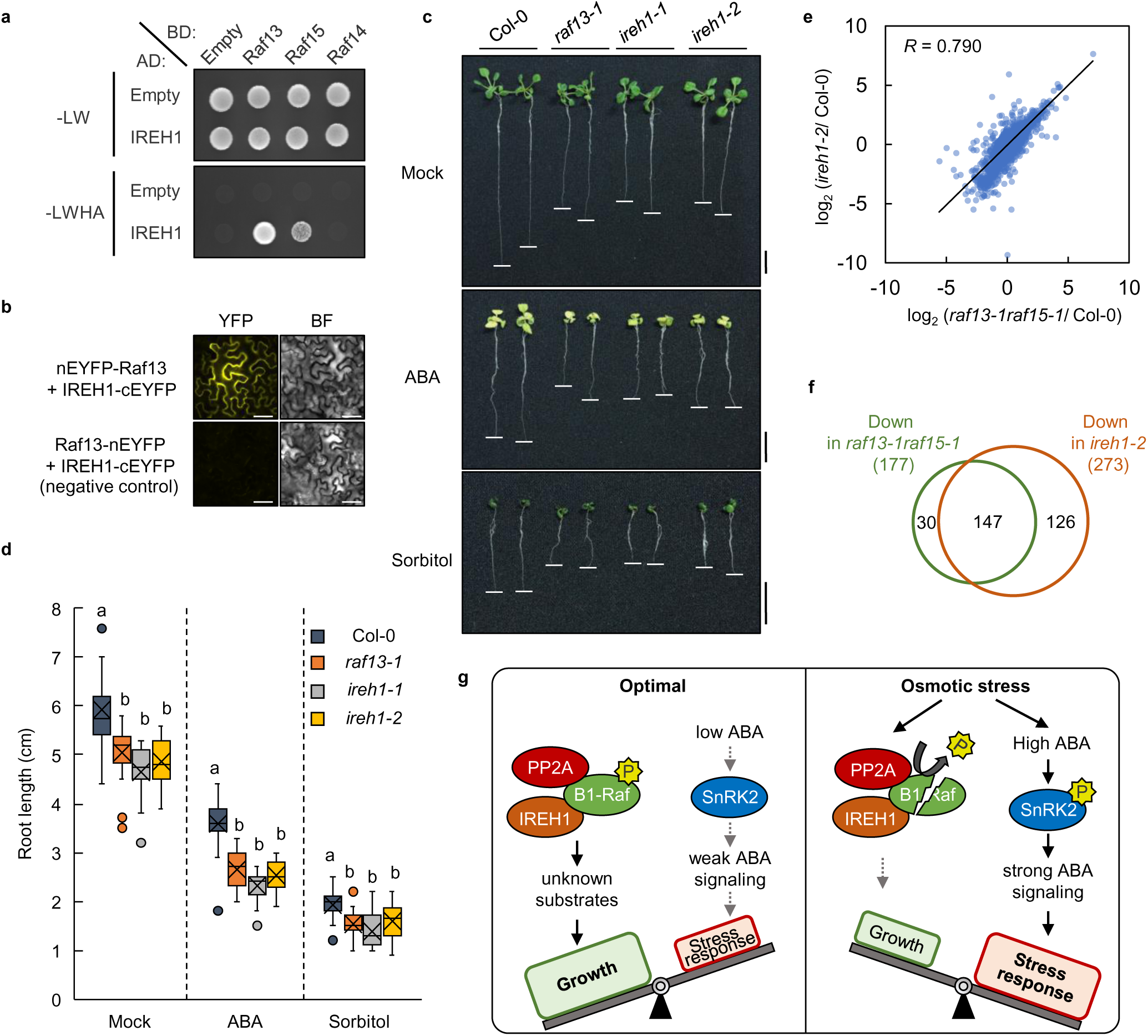
B1-Raf promotes plant growth in cooperation with an AGC protein kinase IREH1. **a**, Y2H assay between B1-Rafs and IREH1. Yeast cells expressing GAL4BD:Raf and GAL4AD:IREH1 fusion proteins were grown on non-selective SD-Leu/ -Trp (-LW) or selective SD-Leu/ -Trp/ -His/ -Ade (-LWHA) media at 30 °C for 5 days. **b,** BiFC assays between Raf13 and IREH1 in *N. benthamiana* epidermal cells. nEYFP and cEYFP represent the N- and C-terminal fragments of the enhanced yellow fluorescent protein (EYFP), respectively. BF indicates brightfield images. Scale bars, 50 μm. **c,d,** Primary root length of 10-day-old *raf13* and *ireh1* mutants under 10 μM ABA or 400 mM sorbitol. Scale bars, 1 cm (**c**). Data are presented as box plots, and different letters indicate significant differences (Tukey’s test, *P* < 0.01, n = 24) (**d**). **e,** Analysis of the correlations between the phosphopeptide abundances in the *raf13-1raf15-1* and *ireh1-2* mutants. The phosphopeptide abundances relative to wild-type in the *raf13-1raf15-1* and *ireh1-2* mutants are shown in the scatter plots together with the corresponding correlation coefficient. Among the 4,062 phosphopeptides detected by LC-MS/MS, 3,946 were included in the analysis in which the average of phosphopeptide abundances in all three genotypes (wild-type, *raf13-1raf15-1* and *ireh1-2*) was not equal to 0. **f,** Venn diagram showing the number of downregulated phosphopeptides in the *raf13-1raf15-1* and *ireh1-2* mutants under normal conditions (two-tailed Student’s *t-*test: *P* < 0.05, fold change < 0.5, compared with wild-type). **g,** Model illustrating the B1-Raf/IREH1/PP2A kinase-phosphatase complex-dependent mechanism for alleviating the trade-off between plant growth and stress responses in Arabidopsis (see also Extended Data Fig. 10).

Cantharidin (CT) is a widely utilized PP2A inhibitor that preferentially inhibits the activities of Ser/Thr protein phosphatases type 1 (PP1) or type 2A (PP2A) ^23^. We tested electrophoretic mobility shifts of Raf13-CFP-HA with or without CT treatment. Raf13-CFP-HA did not show any mobility shifts, but its abundance significantly increased in CT-treated plants (Fig. 4c,d), indicating one or more CT-sensitive protein phosphatases dephosphorylate Raf13 *in vivo*. Furthermore, in amiRNA-based *ATBα/β*-knockdown plants, the hyperosmolarity-induced mobility shift of Raf13-CFP-HA was abolished and was proved as increased protein amounts (Fig. 4e). Note that expressions of both *ATBα* and *ATBβ* mRNA were suppressed by about 55-60% in the two *ATBα/β*-knockdown plants (Fig. 4f).

We then obtained T-DNA insertion mutants for *ATBα* or *ATBβ* (Extended Data Fig. 5b). These mutants displayed ABA-insensitive phenotype in cotyledon greening as well as increased water loss from detached leaves (Fig. 4g,h,i). The mRNA abundances of *ATBα*, but not *ATBβ,* were upregulated after dehydration treatment (Extended Data Fig. 5c). These results indicated that ATBα and ATBβ are bona fide regulators to promote ABA signal transduction along with the acquisition of osmotolerance.

Collectively, these results strongly suggested the involvement of PP2A holoenzymes, which harbor ATBα or ATBβ as regulatory B subunits, in the hyperosmolarity-induced Raf13 dephosphorylation.

### B1-Rafs function cooperatively with an AGC protein kinase IREH1

In addition to PP2A, IP-MS analyses identified Raf15 and AGC kinase IREH1 as Raf13-interacting proteins (Fig. 4a and Supplementary Table 3). We confirmed that IREH1 can interact with Raf13 and Raf15, but not with Raf14, in yeast two-hybrid (Y2H) analyses (Fig. 5a). In addition, bimolecular fluorescence complementation (BiFC) assays demonstrated that IREH1 interacts with Raf13 in cytosol in *N. benthamiana* epidermal cells (Fig. 5b).

We next obtained two *IREH1* T-DNA insertion lines (Extended Data Fig. 6a). Both *ireh1-1* and *ireh1-2* mutants exhibited ABA-hypersensitive phenotype in the greening response (Extended Data Fig. 6b,c), and displayed shorter primary roots which were similar to *raf13-1* (Fig. 5c,d). These results indicated that IREH1 cooperates with B1-Raf for growth promotion and negative regulation of ABA responses.

To gain insight into their signaling pathways, we performed a comparative phosphoproteomic analysis of wild-type, *raf13-1raf15-1* and *ireh1-2* seedlings grown under optimal growth conditions. As a result, phosphoproteomic profiles in *ireh1-2* were significantly correlated with those in *raf13-1raf15-1* (Fig. 5e). Importantly, of the 177 phosphopeptides downregulated in *raf13-1raf15-1*, 83.1% (147/177) overlapped with *ireh1-2* (Fig. 5f and Supplementary Table 4), suggesting that B1-Rafs and IREH1 share phosphorylation targets *in planta*. Notably, gene ontology (GO) analysis reported terms related to growth and development such as cell division, carbon fixation, and flower development from 147 phosphopeptides downregulated in both *raf13-1raf15-1* and *ireh1-2* (Extended Data Fig. 7a and Supplementary Table 5). Three phosphorylation motifs, [-*p*S-P-], [-R-x-x-*p*S-] and [- *p*S-M-], were enriched in the phosphopeptides downregulated in both *raf13-1raf15-1* and *ireh1-2* (Extended Data Fig. 7b). These results suggest that Raf13, Raf15 and/or IREH1 tend to prefer [-*p*S-P-], [-R-x-x-*p*S-] and [-*p*S-M-] motifs or alternatively that Raf13, Raf15 and/or IREH1 regulate kinases that phosphorylate these motifs *in planta*.

Finally, we performed *in vitro* phosphorylation assays using Raf13 and IREH1. Recombinant Raf13 and IREH1 directly phosphorylated kinase-dead forms of IREH1 (K911N) and Raf13 (K546N), respectively, indicating reciprocal phosphorylation between Raf13 and IREH1 (>Extended Data Fig. 8).

## Discussion

Plants potentially have a physiological trade-off between growth and stress responses, yet the full complement of signaling factors and underlying mechanisms are not completely understood. In this study, we hypothesized that Raf13 is involved in such a trade-off in Arabidopsis. Raf13 belongs to B1 subgroup of Raf-like protein kinase family in Arabidopsis, and the B1-Raf subgroup consists of two additional members, Raf14 and Raf15. Although B2-, B3- and B4-Rafs had been well characterized as upstream regulators of SnRK2 in osmotic stress ^9–13^, the functions of B1-Rafs were still unclear.

In this study, our genetic analysis revealed that a function of B1-Rafs is to promote plant growth under non-stress conditions. In this regard, Raf13 functions partially redundantly with Raf15 (Fig. 2a,b,f), and Raf13 and Raf15 may form a heterodimer in vivo (Fig. 4a). Our data also suggest that Raf13 is an active kinase in non-stressed conditions and is destabilized in response to osmotic stress, possibly via rapid dephosphorylation at multiple sites (Fig. 1b-h), resulting in attenuation of growth promotion signaling by B1-Raf in plants. Such a regulation must be helpful to rebalance energy use when plants are exposed to drought stress. It should be noted that remaining dephosphorylation site(s) or other post-translational modification(s) might be required for Raf13 inhibition, because the recombinant MBP-Raf13 with seven Ala substitutions in the N-terminus (Raf13 7A) showed fewer effects on phosphorylation activities than Raf13-CFP-HA immunoprecipitated from dehydration-treated plants (Fig. 1f,g). In addition to this possible mechanism of post-translational inhibition, transcriptional suppression might be another inhibitory system of Raf13 during osmotic stress responses (Extended Data Fig. 9).

By IP-MS analysis we identified an AGC kinase IREH1 that associates with Raf13 (Fig. 4a). The *ireh1* mutants showed similar phenotypic changes to *raf13* mutants, e.g. loss of either IREH1 or Raf13 decreased primary root elongation (Fig. 5c,d), supporting that B1-Raf and IREH1 may be working cooperatively for growth promotion. Furthermore, by IP-MS analysis, we identified ATBα and ATBβ, the two B55-family regulatory B subunits of PP2A, as Raf13-interacting proteins and demonstrated that ATBα/β are required for the hyperosmolarity-induced Raf13 dephosphorylation. That is, *atbα* or *atbβ* mutants displayed opposite phenotypes to the B1-Raf knockout mutants (Fig. 4g,h,i), and hyperosmolality-induced dephosphorylation of Raf13 was diminished by PP2A inhibitor (Fig. 4c,d) or knockdown of *ATBα*/*β* (Fig. 4e,f). The results suggest that PP2A-B55s holoenzymes may directly dephosphorylate Raf13. Given that osmotic stress, but not ABA, induces Raf13 dephosphorylation (Extended Data Fig. 1), together with the negative regulation of responses to exogenous ABA by B1-Rafs or IREH1 are relatively masked under authentic osmotic stress (Fig. 2a,b,c and 5c,d), it is conceivable that the PP2A-mediated inhibition of Raf13 is required to alleviate B1-Raf/ IREH1-dependent growth promotion prior to heightened ABA signaling to occur (Fig. 5g).

The presented data show that Raf13 interacts with Raf15 and IREH1 *in vivo*, and that PP2A holoenzyme(s) may associate with Raf13 via B55-family of PP2A B regulatory subunits. As shown in the IP-MS analysis (Fig. 4a), the kinase-phosphatase complex exists regardless of dehydration stress. Interestingly, functions of this complex were likely to be independent of SnRK2, highlighting the uniqueness of B1-Raf functions, because other B-Raf groups directly regulate SnRK2 to positively regulate stress responses^9–13^. Taken together, our data suggest the following hypothetical model for how B1-Raf and its interacting proteins are involved in growth regulation and stress responses. The Raf13/Raf15/IREH1 kinase complex is in an active state under non-stressed conditions. Upon osmotic stress, the Raf13/Raf15/IREH1-dependent growth signal is alleviated through dephosphorylation-mediated inhibition by PP2A-B55s, allowing for maximal ABA signaling to occur. Our findings uncovered a new layer of the regulatory mechanisms that directly link osmotic stress signal to growth signaling pathway for optimizing the balance between growth and stress responses under fluctuating environment (Fig. 5g).

Further studies will be required to know the mechanism by which Raf13, Raf15 or IREH1 promote plant growth, or negatively regulate ABA responses. IREH1 had been reported to be involved in root morphology because of microtubule misarrangement ^24, 25^. IREH1 is one of 39 members of the AGC protein kinase family in Arabidopsis^26^. In addition to the examples in animals ^27^, it is known that some AGC kinases function in growth signaling pathways in plants, for example, phototropins in blue-light perception and signaling ^28^, PINOID in cellular auxin efflux ^29^, etc. Importantly, among the phosphopeptides downregulated in both *ireh1-2* and *raf13-1raf15-1*, the GO term “cell division” was enriched (Extended Data Fig. 7a), implying that the reduced growth in those mutants might be attributed to some misregulation in cell cycle. Interestingly, the closest orthologs of *Arabidopsis* IREH1 are *Drosophila* and *Xenopus* Greatwall (Gwl), *Homo sapiens* MASTL, budding yeast Rim15, and fission yeast Ppk18, which are essential for mitosis^30–32^. Further analysis will be required to determine whether IREH1 and Raf13/15 play roles in mitotic regulation and are involved in the osmotic stress-dependent cell cycle arrest ^1^.

Intriguingly, in addition to the AGC kinases Gwl/MASTL, the orthologs of ATBα/β (PP2A-B55s) are defined as essential regulators for mitotic cell cycle in animals and fungi. Specifically, Gwl/MASTL promote mitotic entry, whereas PP2A-B55s promote mitotic exit ^32^. PP2A is ubiquitously expressed Ser/Thr protein phosphatase conserved in eukaryotes, and holoenzymes of PP2A function as a heterotrimeric complex, consisting of a catalytic (C), scaffolding (A), and regulatory (B) subunit ^33, 34^. In *Arabidopsis*, there are multiple genes encoding predicted isoforms for each subunit; 3 genes for A, 17 for B, and 5 for C, which can interact in various combinations to exert different regulatory outcomes ^35^. In addition to our findings in ABA/osmotic stress signaling, *Arabidopsis* ATBα and ATBβ have been reported to play roles in plant growth and hormonal responses ^36–39^.

In nature, severe osmotic stress on plants is not likely to occur suddenly, but in most cases, it increases gradually like long-term drought stress ^2^. When the environment is favorable for growth (low osmotic stress), B1-Rafs are highly phosphorylated with higher activity and stability, thereby promoting growth (Extended Data Fig. 10a,b). When plants are exposed to osmotic stress, B1-Raf is dephosphorylated and destabilized to reduce growth, and B2/B3/B4-Rafs are phosphorylated and activated to enhance stress responses. These mechanisms may allow plants to switch from growth to stress responses and survive under the harsh environment (Extended Data Fig. 10a,b). Understanding the molecular mechanisms that sense osmotic stress upstream of B-Rafs will be an important area of future research. Also, functional analyses of substrate candidates for Raf13, Raf15 and/or IREH1, such as those identified in our phosphoproteomic analyses, will help our understanding of how B1-Rafs and IREH1 promote plant growth.

## Methods

### Plant Materials and growth conditions

*Arabidopsis thaliana* ecotype Columbia (Col-0) was used as the wild-type (WT) plants. T-DNA insertion mutant lines, *raf13-1* (SALK_137974C), *raf13-2* (SAIL_104_B07), *raf14-1* (SALK_126902C), *raf15-1* (SALK_119918C), *raf15-2* (SALK_020351C), *atbα* (SALK_095004C), *atbβ* (SALK_062514C), *ireh1-1* (SALK_017861) and *ireh1-2* (SALK_069962C) were obtained from the *Arabidopsis* Biological Resource Center (ABRC) (https://abrc.osu.edu/). The *raf13-1raf15-1* double mutant was generated by crossing *raf13-1* and *raf15-1,* and *raf13-1raf14-1raf15-1* triple mutant (B1-TKO) was generated by crossing *raf14-1* and *raf13-1raf15-1*. The *atark1/2/3* mutant (referred to as B3-TKO in this study) was established as previously described ^9^. Seeds of wild-type, mutants or transgenic plants were sterilized, and sown on 0.8% (w/v) germination agar medium (GM) containing 1% (w/v) sucrose. After vernalization at 4°C in the dark for 4 days, the plants were transferred to a growth chamber with continuous white light illumination of 90 µmol m^-2^ s^-1^ photon flux density and grown for indicated periods at 22°C. For ABA sensitivity test, seeds were sown on GM agar medium with or without indicated concentrations of ABA (Sigma-Aldrich). Germination and greening rates were scored as any seed with root emergence or green cotyledons, respectively.

### Vector constructions

Full-length of the *AtARK1/Raf4* cDNA was previously cloned into pENTR/D-TOPO vector (Thermo Fisher Scientific) ^9^. In addition, full-length of the *Raf13*, *Raf14*, *Raf15*, *ATBα*, *ATBβ* and *IREH1* cDNAs were cloned into pENTR1A vector (Thermo Fisher Scientific) and verified by sequencing. Amino acid substitutions were carried out by site-directed mutagenesis as previously described ^17^. Those cDNAs were transferred into destination vectors, such as pGreen0029-GFP, pGreen0029-HA, R4pGWB 501 ^40^, pEarleyGate 102 ^41^, pSITE-nEYFP-C1 (CD3-1648), pSITE-nEYFP-N1 (CD3-1650), pSITE-cEYFP-N1 (CD3-1651), pGBKT7 and pGADT7 (Takara Bio) by using Gateway LR Clonase II (Thermo Fisher Scientific). For the artificial microRNA to silence the expression of both *ATBα* and *ATBβ*, primer sequences were designed using a Web microRNA designer, WMD3 (http://wmd3.weigelworld.org/cgi-bin/webapp.cgi), and the resulting PCR fragment was cloned into pENTR1A vector to generate pENTR1A harboring *ATBα*/*β-ami*. For the pDONR P4-P1R *Raf13p* or pDONR P4-P1R *UBQ10p* constructs, the putative *Raf13* promoter (2,712 bp upstream sequence) and the *UBQ10* promoter (636 bp upstream sequence) were amplified with specific primers with Gateway attB4/attB1r adaptor sequences and cloned into pDONR P4-P1R vector (Thermo Fisher Scientific) using Gateway BP Clonase II (Thermo Fisher Scientific). For the R4pGWB 501 *Raf13p:GUS* construct, pDONR P4-P1R *Raf13p* and pENTR-gus (Thermo Fisher Scientific) were reacted with R4pGWB 501 ^40^ using Gateway LR Clonase II. In addition, pDONR P4-P1R *UBQ10p* and pENTR1A *ATBα*/*β-ami* were used to establish the R4pGWB 501 *UBQ10p:ATBα*/*β-ami*.

### Transgenic Plants

The pGreen0029 and pEarleyGate 102 were used to express *AtARK1-HA* and *Raf13-CFP-HA*, respectively, as described above. R4pGWB 501 harboring *Raf13p:GUS* was also prepared. Each construct was transformed into *Arabidopsis* wild-type plants with *Agrobacterium tumefaciens* strain GV3101 or GV3101 (pSOUP). Transgenic plants were selected on GM agar medium containing 200 μg/ mL claforan with either 50 μg/mL kanamycin, 25 μg/ mL hygromycin or 10 μg/ mL Basta. For the *35S:Raf13-CFP-HA/ UBQ10p:ATBα*/*β-ami* transgenic plants, the R4pGWB 501 vector harboring *UBQ10p:ATBα*/*β-ami* was transformed into the *35S:Raf13-CFP-HA* transgenic plants and transformants were selected on GM agar medium containing 200 μg/ mL Claforan, 25 μg/ mL hygromycin and 10 μg/ mL Basta.

### Histochemical GUS staining

*Arabidopsis Raf13p:GUS* transgenic seedlings were pretreated with 90% acetone on ice for 15 min, then incubated in GUS staining solution [1 mM 5-bromo-4-chloro-3-indolyl-β-D-glucuronic acid (X-Gluc), 0.5 mM K_3_[Fe(CN)_6_], 0.5 mM K_4_[Fe(CN)_6_], 0.1% (v/v) Triton X-100, 50 mM sodium phosphate (pH 7.2)] in the dark at 37°C for 16 h. The samples were washed and bleached with 70% ethanol. Images of the stained samples were captured using an EPSON GT-X970 scanner (Epson) or using a Leica DMLB fluorescence microscope equipped with a Leica DFC420 camera (Leica).

### Water loss analysis

*Arabidopsis* 7-day-old seedlings grown on 0.8% (w/v) GM agar plates under a 16 h/8 h (light/dark) photoperiod at 22°C were transferred to soil, and the plants were grown under the same conditions for another 3- or 4-weeks. The detached rosette leaves from 4- to 5-week-old plants were placed on weighing dishes and left on the laboratory bench. Fresh weights were monitored as previously described ^5^.

### Measurement of stomatal aperture

Stomatal aperture was measured according to previous studies ^6, 13^. Epidermal strips were peeled from the rosette leaves of 4-week-old *Arabidopsis* seedlings grown in soil and incubated in MES buffer [10 mM MES-KOH (pH 6.2), 10 mM KCl, and 50 µM CaCl_2_] under white light for 3 h to fully open the stomata. Then the strips were transferred to MES buffer containing 10 µM ABA for 2 h. Stomatal apertures were photographed using a BX53 fluorescence microscope (Olympus) and measured by quantifying pore width of stomata using ImageJ software.

### RNA extraction and qRT-PCR

Total RNA was extracted from 2-week-old *Arabidopsis* seedlings as described previously ^5^, and 500 ng of total RNA was used for reverse transcription using ReverTra Ace qPCR RT Master Mix with gDNA Remover (TOYOBO). qRT-PCR analysis was performed using LightCycler 480 SYBR Green I Master (Roche Life Science) with Light Cycler 96 (Roche Life Science). Each transcript was normalized by *GAPDH* and analyzed with three or four biological replicates. The gene-specific primers used for qRT-PCR are listed in Supplementary Table 6.

### Preparation of Recombinant Proteins

pMAL-c5X vectors harboring *SRK2E (K50N)*, *SRK2E (K50N S171A S175A)*, *SRK2G (K33N)* or *AtARK1* cDNAs were previously generated ^9^. In addition, *Raf10, Raf13* and *IREH1* cDNAs were cloned in-frame to pMAL-c5X vector (New England Biolabs) by using In-Fusion HD Cloning Kit (Takara Bio). Amino acid substitutions were introduced into pMAL-c5X Raf13 by site-directed mutagenesis to produce the kinase-dead *Raf13 (K546N)*, phospho-null *Raf13 (7A)* or phospho-mimetic *Raf13 (7D)*. The recombinant proteins were expressed, and affinity purified from *Escherichia coli* strain BL21 (DE3) using Amylose Resin (New England Biolabs) as previously described ^5^.

### In vitro phosphorylation assays

For the kinase assay using bacterially expressed proteins, proteins were mixed with the indicated pair(s) and incubated in a total volume of 10 μL of reaction buffer [50 mM Tris-HCl (pH 7.5), 5 mM MgCl_2_, 5 mM MnCl_2_, 50 µM ATP and 0.037 MBq of [γ-^32^P] ATP (PerkinElmer)] at 30°C for 30 min. To assess the kinase activity of the immunoprecipitated Raf13-CFP-HA protein, approximately 5 g of *Arabidopsis* 2-week-old *35S:Raf13-CFP-HA* transgenic seedlings grown on agar plates were treated with/without dehydration for 30 min on the filter paper, and Raf13-CFP-HA was immunoprecipitated using 20 µL of GFP-selector (NanoTag Biotechnologies, N0310, lot 03210303). The immunoprecipitates were washed three times and then incubated with 2 μg of histone in the same reaction buffer as described above. The protein samples were subsequently separated by a sodium dodecyl sulfate-polyacrylamide gel electrophoresis (SDS-PAGE), and phosphorylation levels were visualized by autoradiography using BAS-5000 (Fujifilm).

### In-gel kinase assays

Crude proteins were extracted from 2-week-old *Arabidopsis* seedlings in extraction buffer [50 mM HEPES-KOH (pH 7.5), 5 mM EDTA, 5 mM EGTA, 1 mM Na_3_VO_4_, 25 mM NaF, 50 mM β-glycerophosphate, 20% (v/v) glycerol and 1% (v/v) protease inhibitor cocktail (Sigma-Aldrich)]. 40 µg of total protein was separated on a 10% SDS-PAGE gel containing histone IIIS as kinase substrate. After electrophoresis, the gel was washed four times at room temperature with wash buffer [25 mM Tris-HCl (pH 7.5), 5 mM NaF, 0.1 mM Na_3_VO_4_, 0.5 mg/mL BSA, 0.1% TritonX-100 and 0.5 mM dithiothreitol (DTT)] and incubated at 4°C for 16 h with three changes of renaturation buffer [25 mM Tris-HCl (pH 7.5), 5 mM NaF, 0.1 mM Na_3_VO_4_ and 1 mM DTT]. The gel was then incubated in reaction buffer [40 mM HEPES-KOH (pH 7.5), 0.1 mM EGTA, 20 mM MgCl_2_ and 1 mM DTT] with 30 µM ATP plus 25 mM [γ-^32^P] ATP for 1 h at 30°C. The reaction was stopped by transferring the gel to wash solution [5% (v/v) trichloroacetic acid (TCA) and 1% (w/v) sodium pyrophosphate], and the gel was washed with the same solution five times. Radioactivity was detected using a BAS -5000 (Fujifilm).

### Western blotting

Proteins were extracted from plants in extraction buffer [20 mM HEPES-KOH (pH 7.5), 100 mM NaCl, 0.1 mM EDTA, 5 mM MgCl_2_, 20% glycerol, 0.5% TritonX-100, 1 mM Na_3_VO_4_, 25 mM NaF and 1% Protease Inhibitor Cocktail (Sigma-Aldrich)] followed by two-step centrifugation at 4°C for 10 min. The supernatants were separated by 8 or 10% SDS-PAGE. After electrophoresis, the proteins were transferred to the polyvinylidene fluoride (PVDF) membrane (0.45 μm, Millipore) and immunoblotted using the primary antibodies, anti-HA [16B12] (BioLegend, 901513, lot B274467, 1: 3,000 dilution) or anti-FLAG (DYKDDDDK) (Wako, 014-22383, lot SAR0168, 1: 3,000 dilution), using Horse anti-mouse IgG-horseradish peroxidase conjugates (Vector Laboratories, PI-2000, lot ZH0513, 1: 3,000 dilution) as the secondary antibody.

### Co-immunoprecipitation (Co-IP) assays

pENTR entry clones harboring *Raf13*, *Raf10*, *AtARK1*, *ATBα* or *ATBβ* were subcloned into pEarleyGate 102 destination vector with C-terminal CFP-HA fusion or into pEarleyGate 202 with N-terminal FLAG epitope tag fusion ^41^. pEarleyGate 100 vector containing *FLAG-SRK2I* was generated as previously described ^5^. Constructs were introduced into *N. benthamiana* leaves with *A. tumefaciens* strain GV3101. After 3 days of incubation, the leaves were ground to a powder in liquid nitrogen, and total proteins were extracted in 10 mL of extraction buffer [20 mM HEPES-KOH (pH 7.5), 100 mM NaCl, 0.1 mM EDTA, 5 mM MgCl_2_, 20% glycerol, 0.5% TritonX-100, 1 mM Na_3_VO_4_, 25 mM NaF and 1% Protease Inhibitor Cocktail (Sigma-Aldrich)], followed by centrifugation at 17,400 × *g* for 10 min at 4°C. The supernatant was incubated with 20 µL of GFP-selector (NanoTag Biotechnologies) for 3 h at 4°C. The immunoprecipitates were washed four times with 1 mL of the extraction buffer without the protease inhibitor cocktail. After washing, the proteins were eluted in 2xSDS sample buffer [125 mM Tris-HCl (pH 6.8), 4% SDS, 10% (w/v) sucrose, 0.004% (w/v) bromophenol blue and 200 mM DTT]. Aliquots were separated by SDS-PAGE and analyzed by western blotting.

### Microscopic analyses of fluorescent proteins

For subcellular localization analysis, pGreen0029-GFP constructs harboring *Raf13, ATBα* or *ATBβ* were introduced into *N. benthamiana* leaves using *A. tumefaciens* strain GV3101 (pSOUP). In addition, for BiFC assays, pSITE-nEYFP-C1 *Raf13*, pSITE-nEYFP-N1 *Raf13* or pSITE-cEYFP-N1 *IREH1* were introduced into *A. tumefaciens* strain GV3101 (p19). The strains were mixed as indicated pairs and infiltrated into *N. benthamiana* leaves. 3 days after infiltration, GFP or complemented YFP fluorescence was observed using a BX53 fluorescence microscope (Olympus).

### Yeast two-hybrid (Y2H) assays

Y2H assays were performed with the MatchMaker GAL4 Two-Hybrid System 3 (Takara Bio). *Saccharomyces cerevisiae* strain AH109 was co-transformed with different pairs of pGADT7 and pGBKT7 harboring IREH1 or B1-Rafs. A single colony for each transformant grown on non-selective SD/ -Leu (L)/ -Trp (W) media was incubated in liquid media, and then tested on selective SD/ -L/ -W/ -His (H)/ -Ade (A) media at 30°C for 5 days.

### Measurement of primary root length

To assess primary root length, seeds were sown on 0.8% (w/v) GM agar plates and grown under continuous white light at 22°C for 4 days. Seedlings were then transferred to 1.0% (w/v) GM agar plates supplemented with/without indicated concentrations of ABA (Sigma-Aldrich) or sorbitol and grown vertically for an additional 7-11 days. Primary root length was measured after the plants were transferred onto the black cloth, and then photographed.

### Sample preparation for IP-MS analyses

For the identification of proteins potentially interacting with Raf13-CFP-HA, 2-week-old *Arabidopsis* wild-type (Col-0) and *35S:Raf13-CFP-HA* transgenic seedlings were grown on GM agar medium for 2 weeks, followed by dehydration stress on filter paper for 30 min. Crude proteins were extracted in 5 mL of extraction buffer [20 mM HEPES-KOH (pH 7.5), 100 mM NaCl, 0.1 mM EDTA, 5 mM MgCl_2_, 20% glycerol, 0.5% TritonX-100, 1 mM Na_3_VO_4_, 25 mM NaF and 1% Protease Inhibitor Cocktail (Sigma-Aldrich)], and then subjected to centrifugation at 4°C for 10 min to remove cellular debris. The supernatant was incubated with 20 µL of GFP-selector (NanoTag Biotechnologies) for 3 h at 4°C. After centrifugation at 4°C for 20 sec, immunoprecipitates were washed four times with 1 mL of the same extraction buffer. Tryptic digestion was performed according to the previous studies ^42, 43^ with some modifications. Briefly, the beads were resuspended in 12 µL of resuspension buffer [100 mM Tris-HCl (pH 9.0), 12 mM sodium *N*-lauroylsarcosinate (SLS), 12 mM sodium deoxycholate (SDC) and 50 mM ammonium bicarbonate], and then 68 µL of digestion buffer [50 mM ammonium bicarbonate, 10 μg/mL Trypsin (Promega) and 1 mM DTT] was added to digest protein samples on beads. After incubation at 37°C for 12 h, peptides were alkylated by adding 20 µL of alkylation buffer [50 mM ammonium bicarbonate and 50 mM 2-iodoacetamide (IAM)]. Surfactants were removed by the addition of 100 µL of ethyl acetate containing 1% trifluoroacetic acid (TFA). The peptides were desalted using C18 Empore disks (GL sciences), dried in a centrifugal concentrator (CC-105, TOMY), and stored at -20°C until the analysis using Orbitrap Exploris 480 (Thermo Fisher Scientific).

### Sample preparation for phosphoproteomic analyses

For the phosphoproteomic analysis under drought treatment, *Arabidopsis* wild-type seeds were sown and grown on GM agar plates under a 16 h/8 h (light/dark) photoperiod at 22°C for 7 days. Seedlings were transferred from GM agar plates to soil, grown in soil for an additional 2 weeks, and then treated with/without water supply for 3, 5 or 9 days. The aerial parts of seedlings were collected and subjected to analysis. For the phosphoproteome to identify the putative Raf13 dephosphorylation sites, *Arabidopsis* wild-type seeds were sown on GM agar plates and grown under continuous white light illumination at 22°C for 2 weeks. The seedlings were then transferred onto the filter paper and subjected to dehydration stress for 30 min. For the phosphoproteome to evaluate B1- Raf- or IREH1-mediated phosphorylation network, 2-week-old *Arabidopsis* wild-type, *raf13-1raf15-1* and *ireh1-2* seedlings grown on GM agar plates under continuous white light at 22°C were collected and subjected to the analysis.

Protein extraction and digestion were performed according to the previous studies ^5, 10^ with some modifications. Briefly, the collected plant tissues were ground to powder in liquid nitrogen, and total proteins were extracted in extraction buffer [100 mM Tris-HCl (pH 9.0), 8 M urea, 2% Phosphatase inhibitor cocktail 2 (Sigma-Aldrich) and 2% Phosphatase inhibitor cocktail 3 (Sigma-Aldrich)], followed by centrifugation at 17,400 × *g* for 20 min at 4°C. The supernatant was then precipitated by the methanol-chloroform precipitation method, and protein pellets were resuspended in digestion buffer [100 mM Tris-HCl (pH 9.0), 12 mM SLS and 12 mM SDC]. Crude extracts containing 400 μg of protein were subjected to the following enzymatic digestion. The solution was reduced with 10 mM DTT for 30 min and then alkylated with 50 mM IAM for 20 min in the dark. After 5-fold dilution with 50 mM ammonium bicarbonate in ultrapure water, proteins were digested with trypsin (Promega; 1:100 (w/w) enzyme-to-protein ratio) overnight at 37°C. Surfactants were removed by addition of an equal volume of ethyl acetate containing 1% TFA, centrifugation at 15,000 × *g* for 2 min, and the aqueous phase was collected for the phosphopeptide enrichment. Phosphopeptides were enriched by hydroxy acid-modified metal oxide chromatography (HAMMOC) method ^44^, and the enriched phosphopeptides were desalted using C18 Empore disks (GL sciences). After desalting, the peptides were dried using a centrifugal concentrator (CC-105, TOMY) and stored at -20°C until use.

### LC-MS/MS-based proteomic analyses

For the phosphoproteomic analysis or IP-MS analysis, the dried peptides were dissolved in 20 μL of 2% (v/v) acetonitrile (ACN) containing 0.1% (v/v) formic acid (FA) and injected into an Easy-nLC 1200 (Thermo Fisher Scientific). Peptides were separated on a C18 nano HPLC capillary column (NTCC-360/75-3, 75 µm ID×15 cm L, Nikkyo Technos) at 300 nL/min by a nonlinear gradient for 140 min for phosphoproteomic analysis, or 40 min for IP-MS analysis. The mobile phase buffer consisted of 0.1% FA in ultrapure water (Buffer A) with an elution buffer of 0.1% FA in 80% ACN (Buffer B). 140-min gradient was run under the following conditions: 0-5 min, B 6%; 5-79 min, B 6-23%; 79-107 min, B 23-35%; 107-125 min, B 35-50%; 125-130 min, B 50-90%; 130-140 min, B 90%, and 40-min gradient was performed as follows: 0-0.5 min, B 6%; 0.5-11.5 min, B 6-23%; 11.5-30.5 min, B 23-40%; 30.5-35 min, B 40-50%; 35-35.5 min, B 50-90%; 35.5-40 min, B 90%. The Easy-nLC 1200 was coupled to an Exploris 480 quadrupole-Orbitrap mass spectrometer (Thermo Fisher Scientific) with an FAIMS Pro high-field asymmetric waveform aerodynamic ion mobility spectrometry (FAIMS) device (Thermo Fisher Scientific). The mass spectrometer was operated in the data-dependent acquisition (DDA), positive ion mode in which MS1 spectra (375-1500 *m/z* for phosphoproteomic analysis or 350-1200 *m/z* for IP-MS, with a resolution of 60,000) were followed by MS2 spectra (over 120 *m/z* with the resolution of 30,000 for phosphoproteomics or 15,000 for IP-MS). For FAIMS, three conditions of compensation voltage (CV) values (-40/-50/-60V, -40/-60V, - 40/-70V, or -50/-70V) were used with the following settings (standard resolution; inner temperature (IT) 100°C / outer temperature (OT) 100°C).

Peptide/Protein identification and MS1-based label-free quantification were carried out using Proteome Discoverer 2.5 (PD2.5) (Thermo Fisher Scientific) or Skyline version 22.2 (MacCoss lab software). MS2 spectra were searched with SEQUEST HT against the *Arabidopsis* protein database (Araport11_genes.201606.pep.fasta). The SEQUEST search parameters were set as follows: digestion enzyme trypsin, maximum missed cleavages 2, peptide length 6-144, precursor mass tolerance 10 ppm, fragment mass tolerance 0.02 Da, static modification: cysteine (C) carbamidomethylation, variable modification: methionine (M) oxidation/ N-terminal acetylation/ serine (S) threonine (T) tyrosine (Y) phosphorylation, maximum variable modifications 3. Peptide validation was performed using the Percolator, and only high-confidence peptides with a false discovery rate (FDR) < 1% were used for protein inference and quantification. Site localization probability of phosphorylation was calculated with IMP-ptmRS node implemented in PD2.5, and 75% was used as the cut- off for localization of phosphorylation site(s).

### Accession numbers

Sequence data for the genes described in this article are avairable at TAIR (http://www.arabidopsis.org/) under the following accession numbers: Raf13 (AT2G31010), Raf14 (AT2G42640), Raf15 (AT3G58640), Raf4 (AT1G18160), Raf5 (AT1G73660), Raf6 (AT4G24480), Raf10 (AT5G49470), ATBα (AT1G51690), ATBβ (AT1G17720), IREH1 (AT3G17850).

### Data availability

LC-MS raw data have been deposited in Japan Proteome Standard Repository/Database (jPOST; https://repository.jpostdb.org/preview/12483870706419549fc20cf, access key; 1792).

## Acknowledgements

We thank Dr. Yoichi Sakata (Tokyo University of Agriculture, Japan) for providing the B3-TKO (*atark1/2/3*) mutant and Dr. Tsuyoshi Nakagawa (Shimane University, Japan) for providing the R4pGWB 501 vector. We also thank Dr. Jeffrey C. Anderson (OSU, USA) for comments about and proofreading of this manuscript. We are grateful to the ABRC for providing Arabidopsis T-DNA insertional mutants.

---

**Extended Data Fig. 1.**
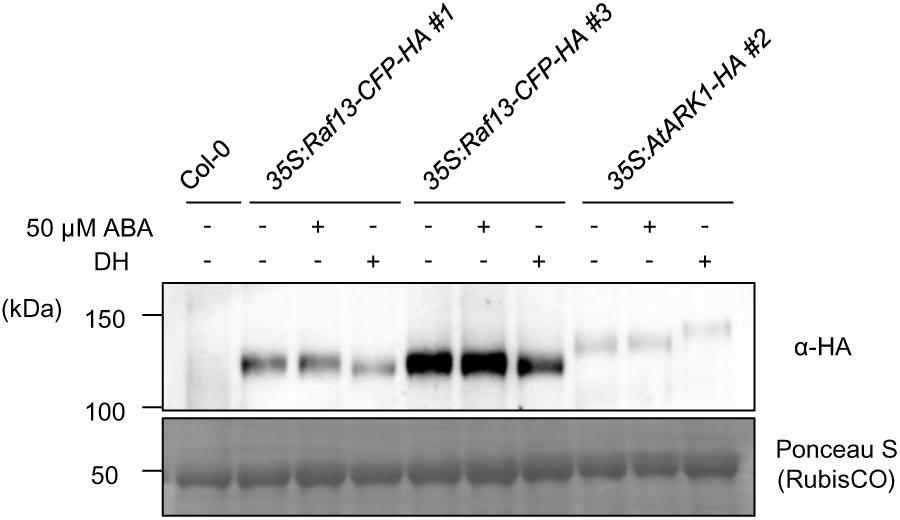
Osmotic stress, but not ABA, induces Raf13 dephosphorylation. Band shifts of Raf13- CFP-HA or AtARK1-HA protein after ABA or dehydration treatment. Crude protein was extracted from 2-week- old wild-type (Col-0), *35S:Raf13-CFP-HA* or *35S:AtARK1-HA* transgenic seedlings after treatment with 50 μM ABA or dehydration (DH) treatment for 30 min.

**Extended Data Fig. 2.**
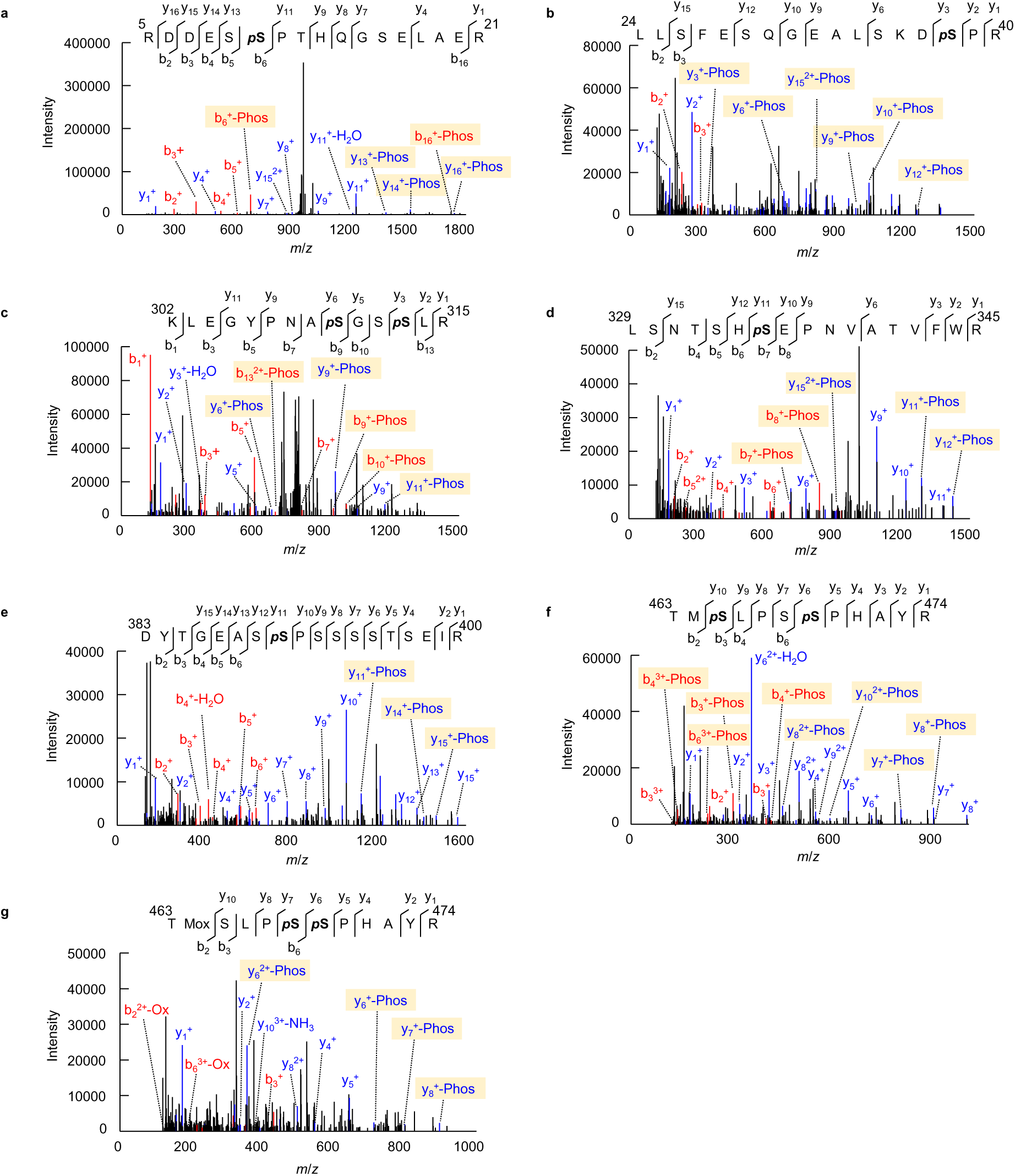
The MS/MS spectra of Raf13 phosphopeptides detected by LC-MS/MS. The MS/MS spectrum shows the phosphopeptide containing the phosphoserine(s); Ser10 (**a**), Ser38 (**b**), Ser310 and Ser313 (**c**), Ser335 (**d**), Ser390 (**e**), Ser465 and Ser469 (**f**), and Ser468 and Ser469 (**g**), in Raf13. Mox indicates methionine oxidation.

**Extended Data Fig. 3.**
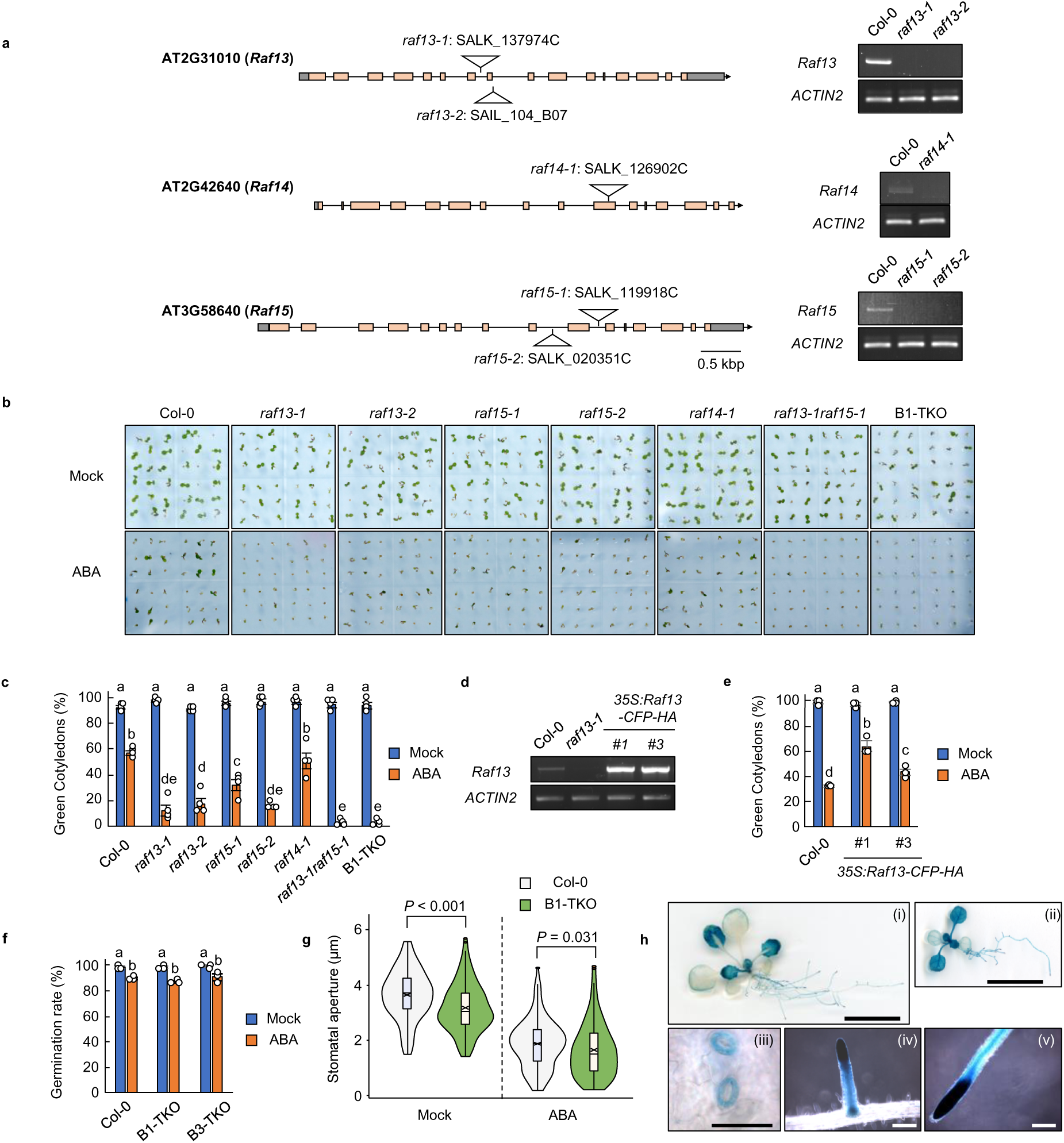
Isolation and characterization of mutants for group B1 Raf-like protein kinases. **a**, Schematic representation of the genomic DNA sequence showing the positions of the T-DNA insertions in *Raf13*, *Raf14* or *Raf15* genes. Light orange boxes and lines indicate exons and introns, respectively. RT-PCR images show the loss of transcripts. *ACTIN2* was used as a positive control. **b,c,** Cotyledon greening rates of wild-type (Col-0) and B1-Raf mutants with or without 0.25 μM ABA for 3 days. Data are means ± SE (n = 4), and different letters indicate significant differences (Tukey’s test, *P* < 0.05). Each replicate contains 36 seeds. **d,e,** RT-PCR analysis of *Raf13* transcript levels (**d**), and the cotyledon greening rates of wild-type and *35S:Raf13-CFP-HA* (*#1* and *#3*) with or without 0.75 µM ABA for 7 days (**e**). Data are means ± SE (n = 3). Each replicate contains 54 seeds. Different letters indicate significant differences (Tukey’s test, *P* < 0.05). **f,** Germination rates of wild-type, B1-TKO and B3-TKO with or without 2 μM ABA for 2 days. Data are mean ± SE (n = 3), and different letters indicate significant differences (Tukey’s test, *P* < 0.05). Each replicate contains 36 seeds. **g,** Violin plots showing the stomatal aperture of 4-week-old wild-type and B1-TKO treated with/without 10 μM ABA for 2 h. The box limits represent the first and the third quartiles with the medians and the means marked as horizontal lines and cross marks, respectively. Significance was determined using two-tailed Student’s *t-*test. (Col-0 Mock, n = 114; B1-TKO Mock, n =127; Col-0 ABA, n = 144; B1-TKO ABA, n = 164, from the six individual leaves). h, The GUS staining of Arabidopsis *Raf13p:GUS* plants. Pictures show the whole seedlings (i and ii), guard cells (iii), lateral root (iv) and primary root (v); 14-day-old (i, iii, iv and v) and 10-day-old (ii); scale bars, 1 cm (i and ii), 50 μm (iii) and 200 μm (iv and v).

**Extended Data Fig. 4.**
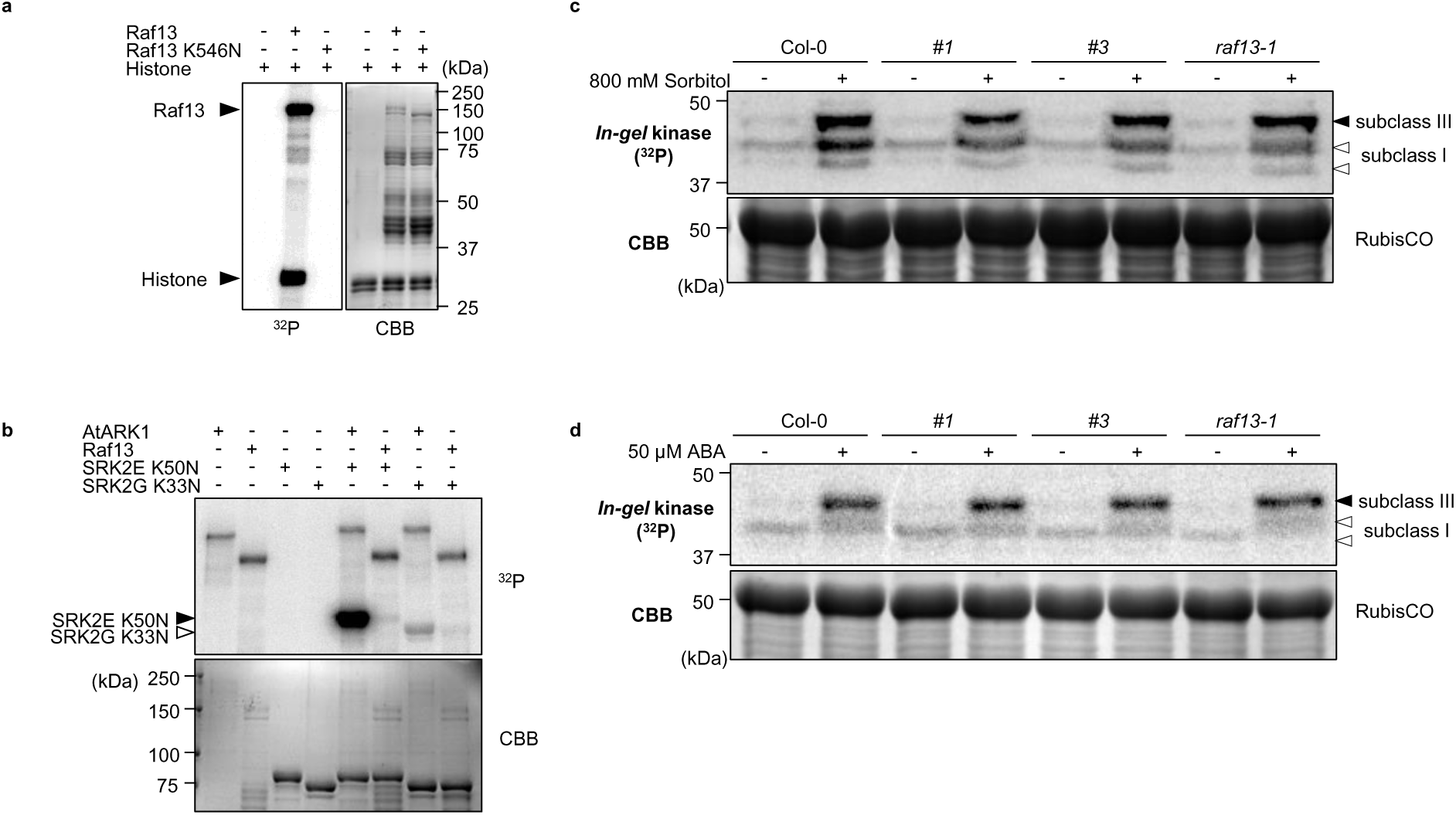
Raf13 functions independently of SnRK2 regulation. **a**, *In vitro* phosphorylation assay showing that Raf13 has a protein kinase activity. Raf13 and kinase-dead Raf13 (K546N) were purified as recombinant MBP fusions. **b,** *In vitro* phosphorylation of kinase-dead forms of subclass III SnRK2 (SRK2E K50N) or subclass I SnRK2 (SRK2G K33N) by AtARK1 or Raf13. All proteins were purified as recombinant MBP fusions. **c,d,** *In-gel* kinase assays of proteins extracted from 2-week-old seedlings of wild-type (Col-0) and two independent *35S:Raf13-CFP-HA* transgenic plants (*#1* and *#3*) treated with 800 mM sorbitol (**c**) or 50 µM ABA (**d**) for 30 min using histone IIIS as substrate. Black arrows and open arrows indicate the position of subclass III SnRK2s and subclass I SnRK2s, respectively. Autoradiography (^32^P) and Coomassie brilliant blue (CBB) staining in **a,b,c,d** show protein phosphorylation and loading, respectively.

**Extended Data Fig. 5.**
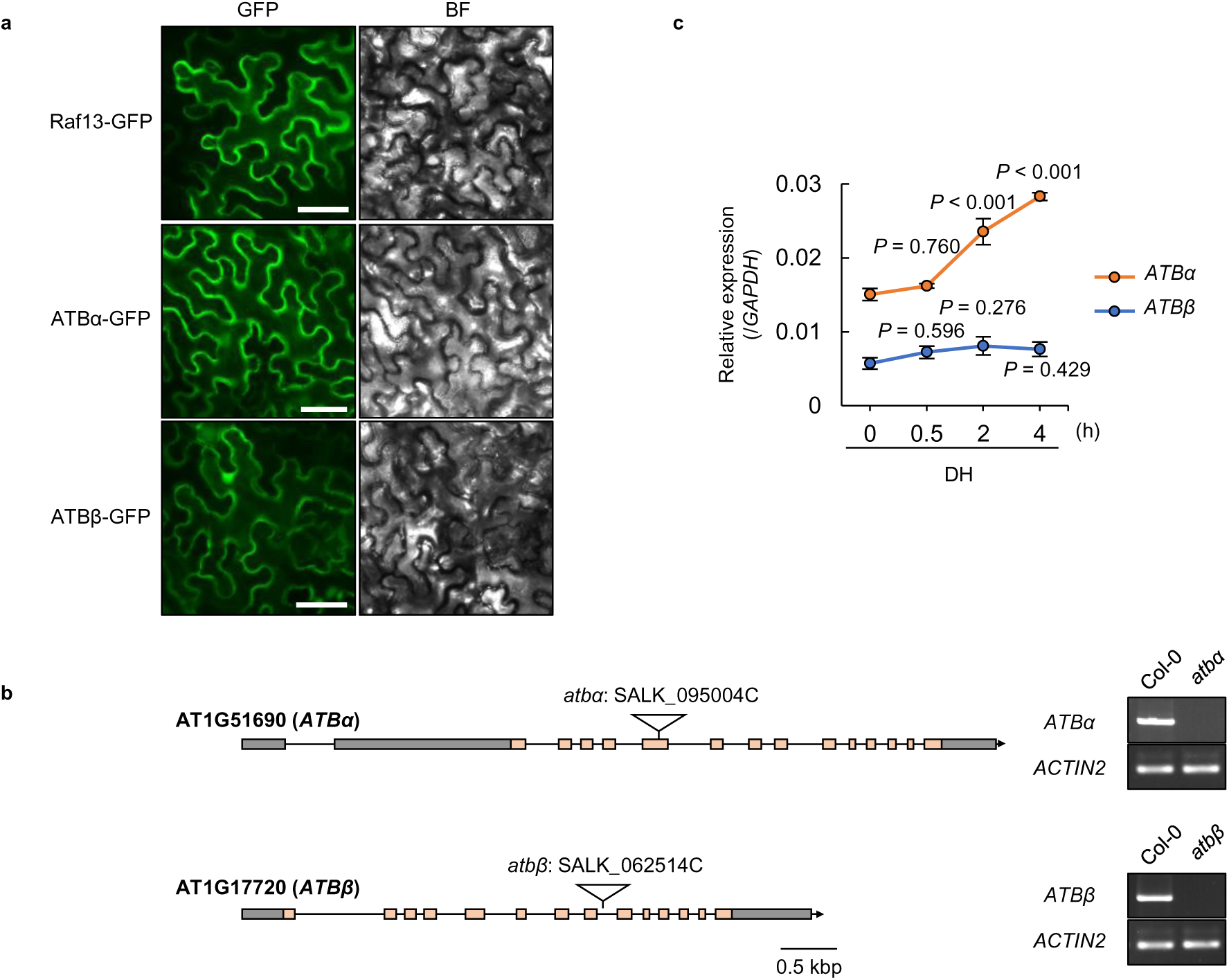
Isolation and characterization of PP2A B subunit ATBα/β mutants. **a**, Subcellular localization of Raf13-GFP, ATBα-GFP or ATBβ-GFP transiently expressed in *N. benthamiana* epidermal cells. BF indicates brightfield images. Scale bars, 50 μm. **b,** Schematic representation of the genomic DNA sequence indicating the locations of the T-DNA insertions in *ATBα* or *ATBβ*. Light orange boxes and lines indicate exons and introns, respectively. Loss of transcripts in *atbα* or *atbβ* mutants was confirmed by RT-PCR. *ACTIN2* was used as a positive control. **c,** Relative transcript levels of *ATBα* or *ATBβ* in 2-week-old Arabidopsis wild-type plants treated with/without dehydration (DH) were measured by qRT-PCR. *GAPDH* was used as an internal control. Data are means ± SE (n = 3), and significance was determined by Dunnett’s test between relative expression at 0 h and that of at each time point.

**Extended Data Fig. 6.**
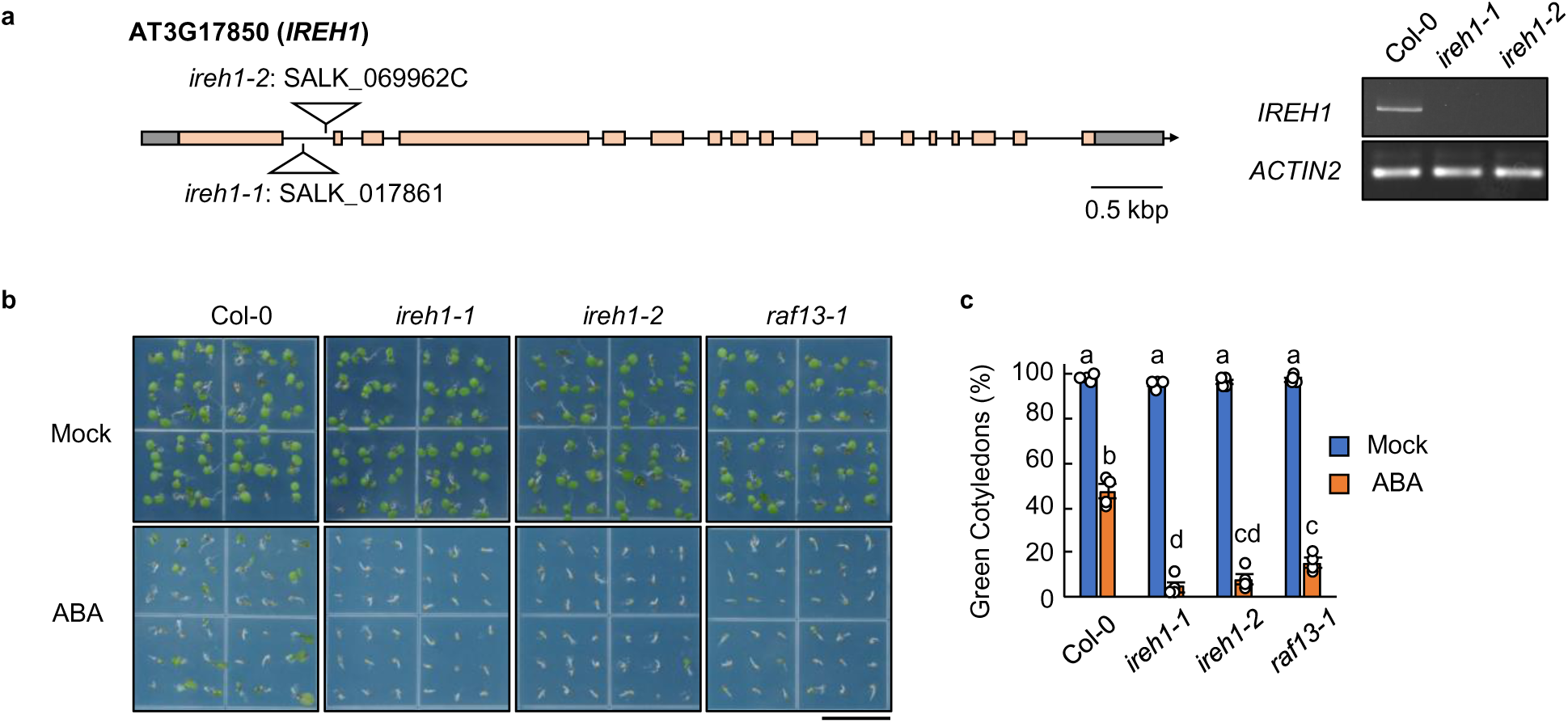
Isolation and characterization of *ireh1* mutants. **a**, Schematic representation of the genomic DNA sequence showing the locations of the T-DNA insertions in *IREH1*. Light orange boxes and lines indicate exons and introns, respectively. The loss of *IREH1* transcripts in *ireh1-1* and *ireh1-2* mutants was confirmed by RT-PCR. *ACTIN2* was used as a positive control. **b,c,** Cotyledon greening of wild-type (Col-0), *ireh1-1*, *ireh1-2* and *raf13-1* plants with/without 0.5 μM ABA for 4 days (**b**), and data are presented as means ± SE (n = 4) (**c**). Different letters indicate significant differences (Tukey’s test, *P* < 0.05). Each replicate contains 54 seeds. Scale bar, 1 cm.

**Extended Data Fig. 7.**
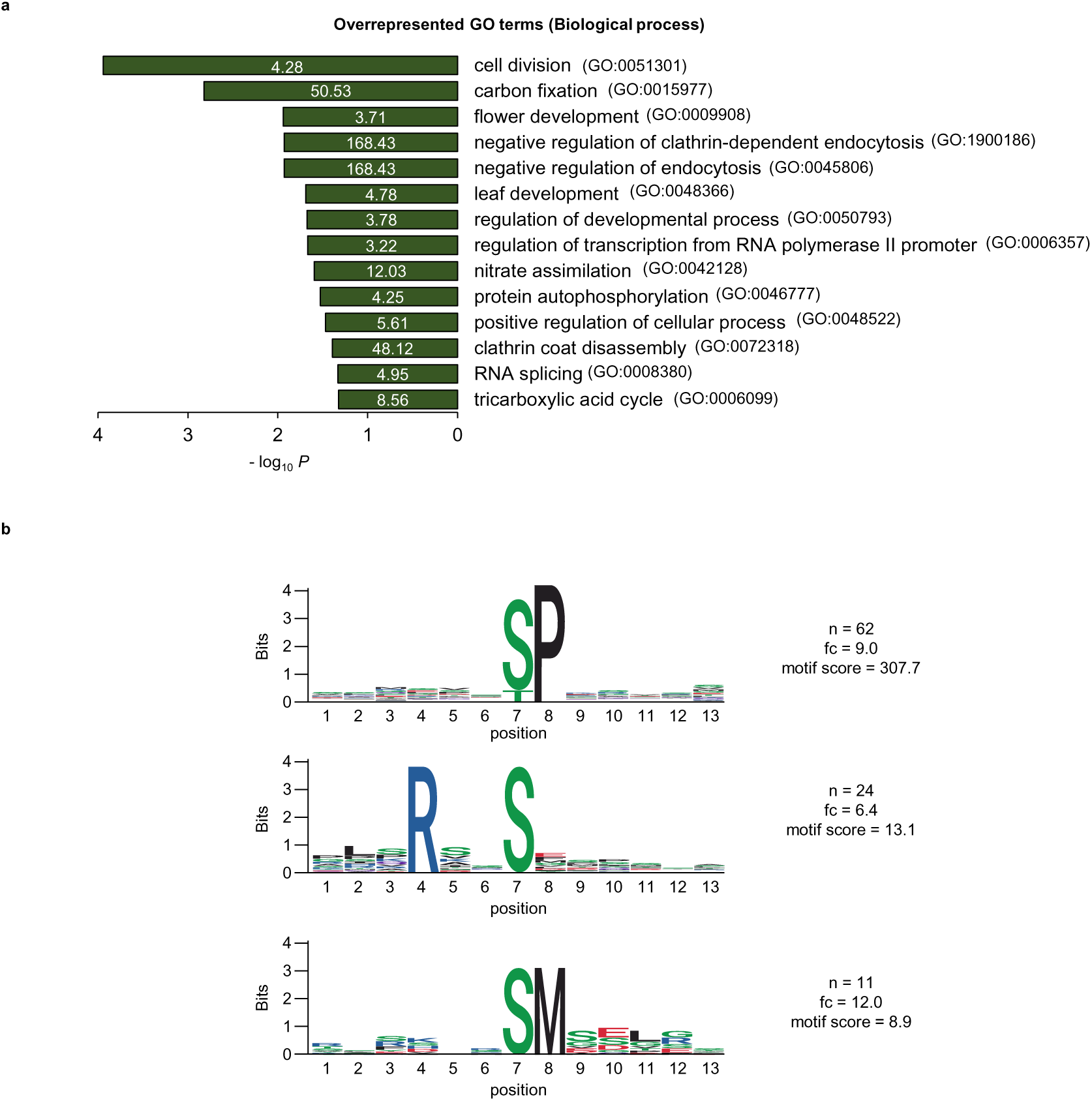
Comparative phosphoproteomic analyses of wild-type, *raf13-1raf15-1* and *ireh1-2*. **a**, Enriched GO terms for 147 phosphopeptides downregulated in both *raf13-1raf15-1* and *ireh1-2* mutants evaluated by DAVID program. The numbers in the bars represent the fold enrichment that were determined by the number of query genes divided by expected number of genes in each GO term. **b,** Motif analysis of phosphopeptides downregulated in both *raf13-1raf15-1* and *ireh1-2* mutants. 13 amino acids around phosphorylated serine (S)/ threonine (T)/ tyrosine (Y) residues were extracted from identified phosphopeptide sequences and analyzed using the Motif-X algorithm implemented in the rmotifx R package.

**Extended Data Fig. 8.**
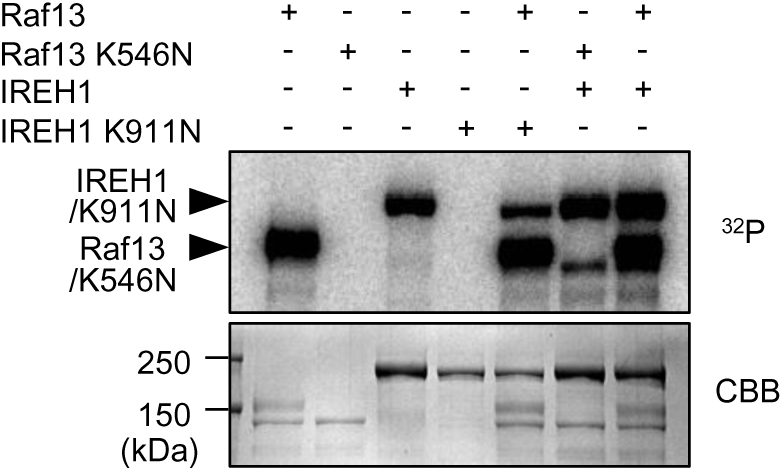
*In vitro* phosphorylation between Raf13 and IREH1. *In vitro* phosphorylation assay using kinase-dead forms of MBP-Raf13 (Raf13 K546N) or MBP-IREH1 (IREH1 K911N). Each kinase-dead form was co-incubated with an active MBP-IREH1 or MBP-Raf13 kinase as indicated. Autoradiography (^32^P) and CBB staining (CBB) show protein phosphorylation and loading, respectively.

**Extended Data Fig. 9.**
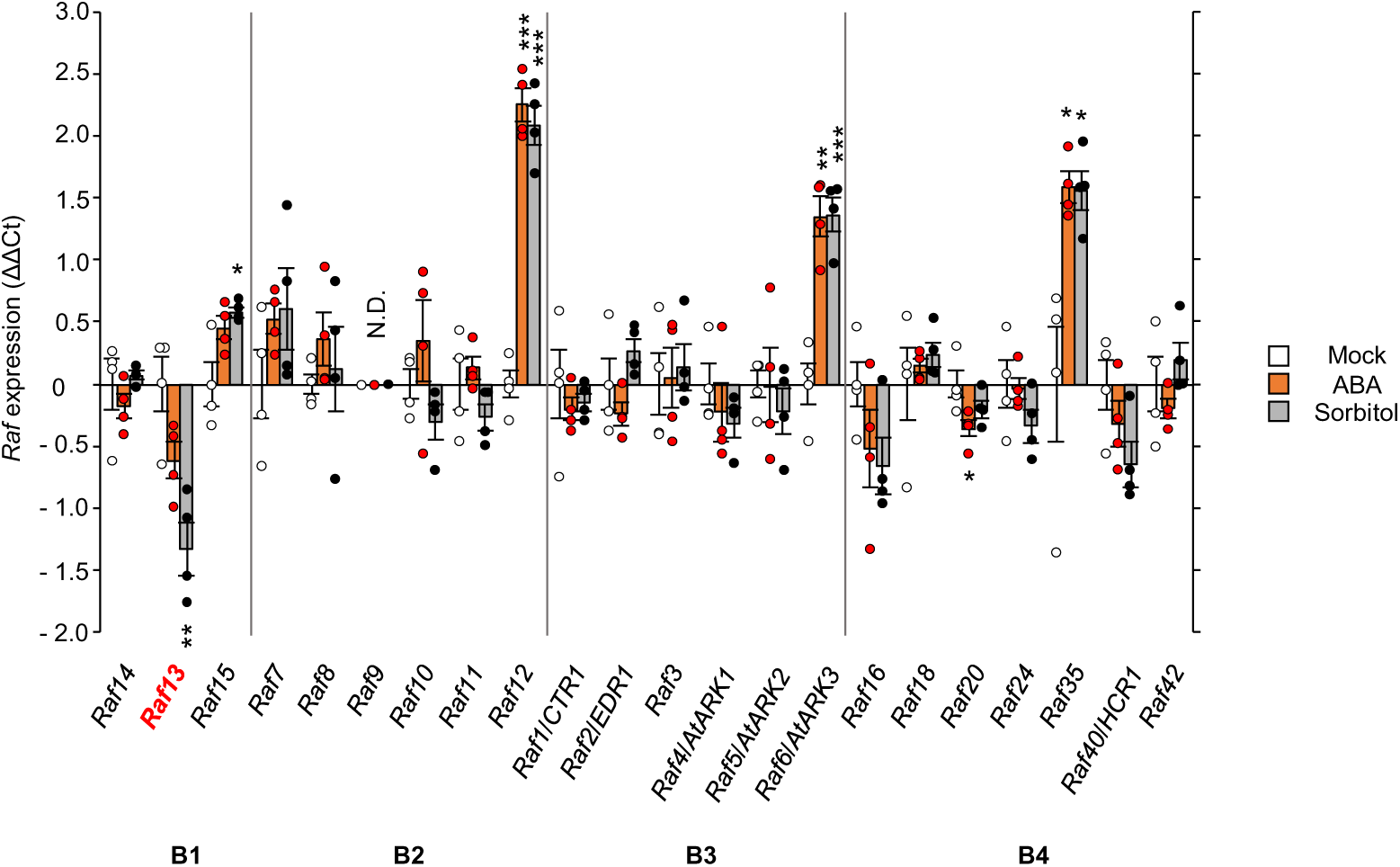
Gene expression levels of Arabidopsis group B Raf-like protein kinases. Arabidopsis 2-week-old wild-typeseedlings (Col-0) were treated with water (Mock), 10 μM ABA or 300 mM Sorbitol for 3 h, and gene expression levels were measured by qRT-PCR. Relative transcript levels are expressed as ΔΔ Ct. ΔΔ Ct = Δ Ct (Mock) – Δ Ct (ABA or Sorbitol), where Δ Ct = Ct (*Raf*) – Ct (*GAPDH*). Data are mean ± SE from four independent biological replicates. Asterisks indicate significant differences (\*\*\**P* < 0.001, \*\**P* < 0.01, \**P* < 0.05, two-tailed Student’s *t* test). N.D., not detected.

**Extended Data Fig. 10.**
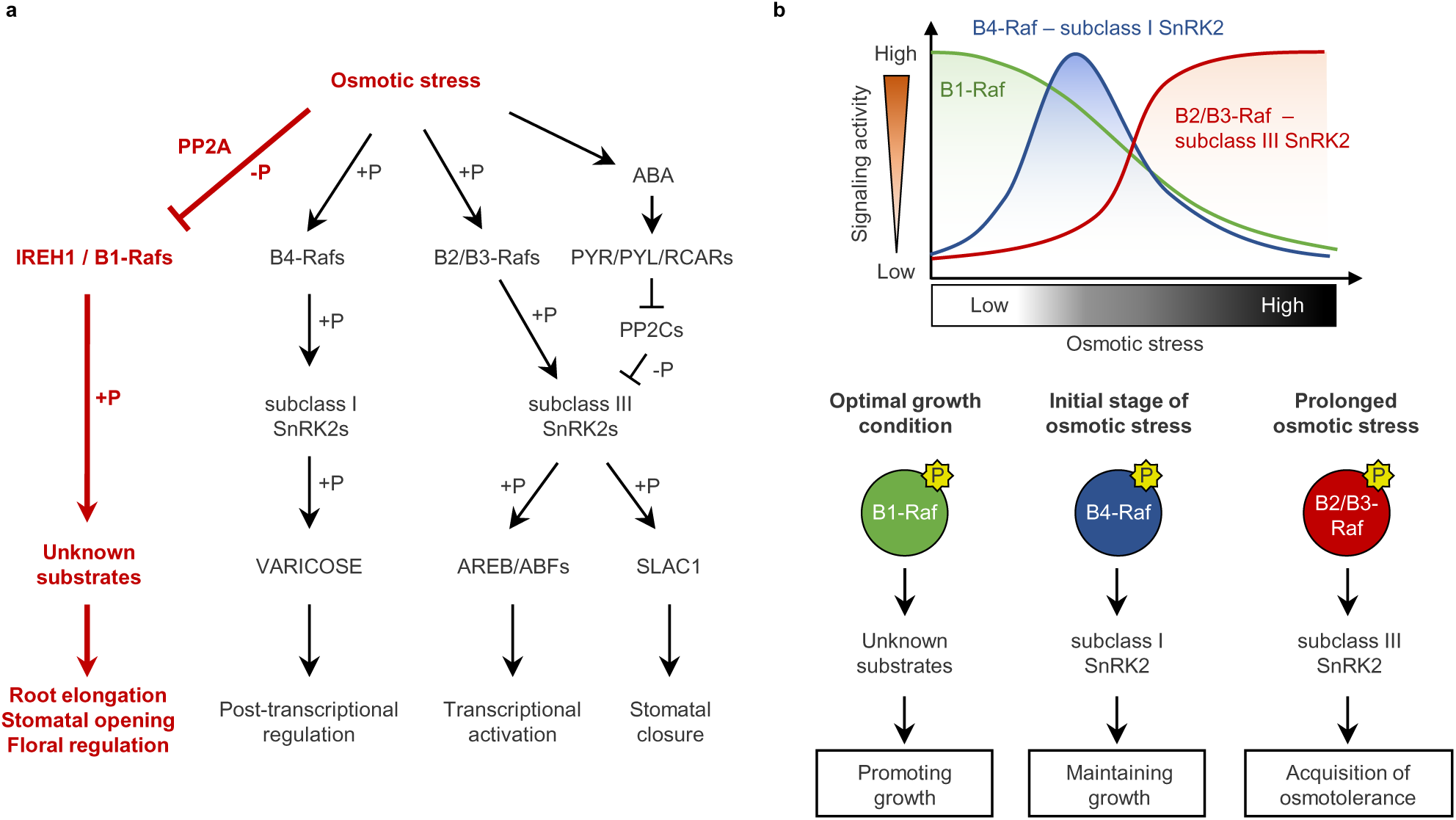
Group B Raf-like protein kinases regulate the balance between growth and osmotic stress responses in *Arabidopsis*. **a**, Model of group B Raf-dependent signaling in response to osmotic stress. **b,** Graphical illustration showing the activation of the B-Raf signaling at different phases of drought stress.

**Supplementary Table 1. Phosphoproteomic analyses of drought-stressed Arabidopsis wild-type plants.** Phosphopeptide abundances of group B-Rafs, SnRK2s or SnRK2 substrates in the aerial parts of soil-grown Arabidopsis wild-type (Col-0) seedlings with/without water supply for the indicated time periods. Data are obtained from three independent biological replicates. Phosphopeptide abundances were manually inspected and quantified based on the extracted ion chromatogram (XIC) peak area using Skyline software (ver. 22.2).

**Supplementary Table 2. Phosphoproteomic analyses of Arabidopsis wild-type seedlings in response to dehydration treatment.** Phosphopeptide abundances in 2-week-old Arabidopsis wild-type (Col-0) seedlings treated with/without dehydration on filter paper for 30 min. Data were obtained from three independent biological replicates. The phosphopeptides were identified and quantified based on the peak height of the extracted ion chromatogram (XIC) using Proteome Discoverer 2.5 (PD2.5). Missing values were treated as 0. “TRUE” indicates the phosphopeptides with more than 2-fold change (up-regulated by dehydration) and with less than 1/2-fold change (down-regulated by dehydration) with P < 0.05 (two-tailed Student’s t test).

**Supplementary Table 3. List of Raf13-CFP-HA-interacting proteins identified by IP-MS analysis.** Immunoprecipitation (IP) was performed using GFP agarose on protein extracts from agar-plate-grown Arabidopsis wild-type (Col-0) or 35S:Raf13-CFP-HA transgenic seedlings with/without dehydration (DH) treatment for 30 min, and the immunoprecipitates were analyzed by LC-MS/MS. The peptide spectrum match (PSM) scores in the two independent biological replicates were shown. The cut-off was made with the PSM scores greater than or equal to 10 in 35S:Raf13-CFP-HA in either Mock or DH conditions and less than 6 for wild type in both Mock and DH conditions.

**Supplementary Table 4. Phosphoproteomic analyses of raf13-1raf15-1 and ireh1-2 mutants under optimal growth conditions.** Phosphopeptide abundances in 2-week-old Arabidopsis wild-type (Col-0), *raf13-1raf15-1* or *ireh1-2* seedlings under optimal growth conditions. The data were obtained from three independent biological replicates. Phosphopeptides were identified and quantified based on the peak height of the extracted ion chromatogram (XIC) using Proteome Discoverer 2.5 (PD2.5). Missing values were treated as 0. “TRUE” indicates the phosphopeptides with less than 1/2-fold change compared with wild-type with *P* < 0.05 (two-tailed Student*’*s *t-* test).

**Supplementary Table 5. List of GO terms from phosphopeptides downregulated in both *raf13-1raf15-1* and *ireh1-2*.** GO terms from phosphopeptides downregulated in both *raf13-1raf15-1* and *ireh1-2* mutants as compared with wild-type. GO terms were evaluated using the DAVID program.

**Supplementary Table 6. Primer sequences used in this study.**

